# Endothelial Resolution of Inflammation is Delayed Following JAK-driven but not NFκB-dependent Activation

**DOI:** 10.1101/2021.06.18.449043

**Authors:** Nicole M Valenzuela

## Abstract

Blood endothelial cells actively regulate egress of leukocytes into peripheral tissues in response to inflammatory insult. The resolution of inflammation is critical for healing and return to homeostasis, but the timing and mechanisms involved in return to a non-inflamed state are not well-understood. We examined vascular endothelial activation comparing NFκB-driven TNFα and JAK/STAT-mediated IFNγ. Pro-adhesive gene expression, phenotype and secretome of human endothelial cells from 6 vascular beds were measured under chronic cytokine stimulation, and after short-term cytokine priming followed by withdrawal. The majority of inducible TNFα effectors require continuous exposure for reinforcement of the altered phenotype. NFκB and target genes are quickly down-regulated in the absence of cytokine. In contrast, the consequences of even short exposure to IFNγ are long-lasting and broad, with sustained elevation of adhesion molecules and chemokines up to 48hr later. JAK/STAT and interferon response factor expression are likewise durable, dependent on new transcription and autonomous of continuous IFNγ. Finally, intact persistent STAT expression and JAK signaling in the endothelium is required to maintain a pro-adhesive phenotype after IFNγ withdrawal, which could be prevented by the JAK1/2 inhibitor ruxolitinib. Our results reveal a sustained JAK-dependent perturbation of endothelial function after exposure to IFNγ, but not after NFκB-driven inflammation.

## Introduction

Vascular inflammation underlies numerous cardiovascular diseases, vasculitis, and organ transplant rejection. Endothelial cells (EC) form a selectively permeable barrier between the blood and the tissue. Vascular endothelium actively regulates access of the cellular immune compartment to interstitial spaces, and is exquisitely sensitive and responsive to myriad inflammatory stimuli to facilitate peripheral immune responses. The best-studied immune function of endothelial cells is upregulation of adhesion molecules and chemoattractants to promote infiltration of immune cells at sites of inflammation. Endothelial cells upregulate adhesion molecules that facilitate tethering, firm adherence, and extravasation of leukocytes through the vessel wall, and secrete chemokines to enhance and guide adherence ^1^. Although there is some redundancy, the pattern of inducible adhesion molecules and chemokines can be context and stimulus dependent, with both convergent and distinct intracellular signaling and functional effects. Moreover, the profile of adhesion molecule and chemoattractants influences the selectivity of leukocyte subsets called to sites of inflammation.

Resolution is a key phase of inflammation, leading to reduction in the influx of immune cells at local sites, followed by restoration of normal function. Failure to attenuate this process leads to a detrimental, chronic response and maladaptive immunity ^2,3^. Much has been studied on the initiation of endothelial inflammation, but less is known about the kinetics and mechanisms sustaining inflammation or those actively facilitating the return to a non-inflamed state. Moreover, such pro-resolving mediators may be exogenous or cell-intrinsic, and likely differ across cell types and signaling pathways.

The adaptive cytokines TNFα and IFNγ drive many autoimmune and autoinflammatory diseases. These cytokines trigger highly divergent signaling and transcriptional programs. We recently demonstrated that the antigen presentation capacity of endothelial cells was highly durable following IFNγ but not TNFα activation ^4^. Here, we hypothesized that the dynamics and resolution of endothelial cell pro-adhesive phenotypes might also vary, and sought to dissect the differential intracellular signaling that contributed to a persistent vs. resolvable inflammatory state. Using human endothelial cells, we tested the response after short exposure to TNFα or IFNγ, and investigated the contribution of the predominant nonredundant signaling cascades mediated by NFκB and JAK/STAT pathways. Our results show that TNFα withdrawal results in a prompt contraction of NFκB signaling and downstream adhesion molecule and chemokine expression, whereas IFNγ has a lasting impact on endothelial cell dysfunction, which was dependent on JAK1/2. Our results suggest that NFκB-controlled vascular inflammation is more sensitive to context and ongoing stimulus, compared with JAK/STAT-driven endothelial responses. These findings hold important implications for the capacity of endothelium to promote inflammation resolution following acute activation. Moreover, we demonstrate a role for JAK/STAT in endothelial dysfunction and prolonged vascular inflammation, and propose that JAKinibs may be an untapped therapeutic avenue effective in blunting the endothelial cell inflammatory response.

## Results

HMEC-1 human endothelial cells were treated with TNFα or IFNγ for 1hr, 3hr, 6hr, 18hr and 24hr. Cytokine dosing was based on preliminary titration studies testing TNFα from 6.25-50ng/mL and IFNγ from 31.25-250U/mL (**Supplemental Figure 1**). The heat map (**Figure 1**) shows a temporal analysis of TNFα-induced and IFNγ-induced adhesion molecules and chemokines at the mRNA level.

**Figure 1.**
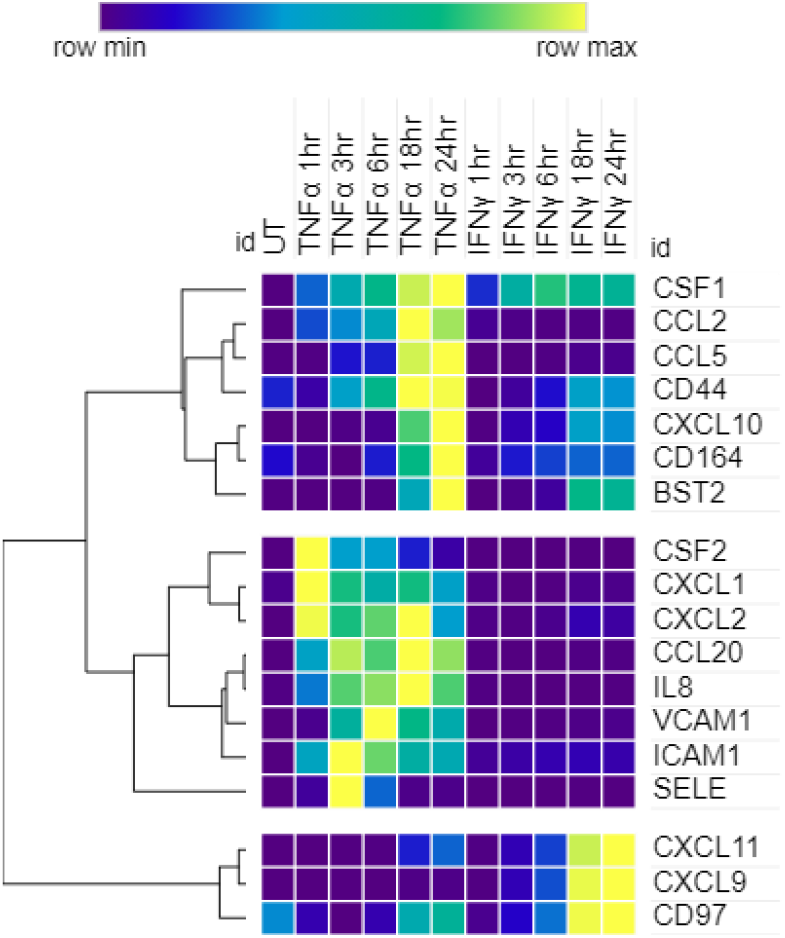
Time course of TNFα and IFNγ stimulated endothelial cell adhesion molecule and chemokine transcripts. HMEC-1 were stimulated with TNFα (20ng/mL) or IFNγ (200U/mL) for 1hr, 3hr, 6hr, 18hr, and 24hr. Adhesion molecule and chemokine mRNA was measured in cell lysates by Nanostring. One representative experiment is shown. Heat map shows relative normalized mRNA counts for each gene.

### Time course of TNFα-induced adhesion molecules

We next measured the TNFα-induced kinetics of pro-adhesive genes in primary endothelium, testing mRNA expression in a public dataset of HUVEC treated for 1-6hr (GSE27870), and protein expression in HAEC treated for 3-24hr.

E-selectin mRNA was not detected in untreated EC. *SELE* transcripts rose by 1hr, peaked at 3hr, and returned to baseline by 18hr of TNFα stimulation (**Figure 2a, 2b**). Similarly, E-selectin protein became detectable as early as 3hr after TNFα exposure, peaking at 6hr (50.92±25.65-fold) and then declining to baseline levels by 18-24hr (9.67±4.68-fold) and remaining undetectable through 40hr (**Figure 2c**).

**Figure 2.**
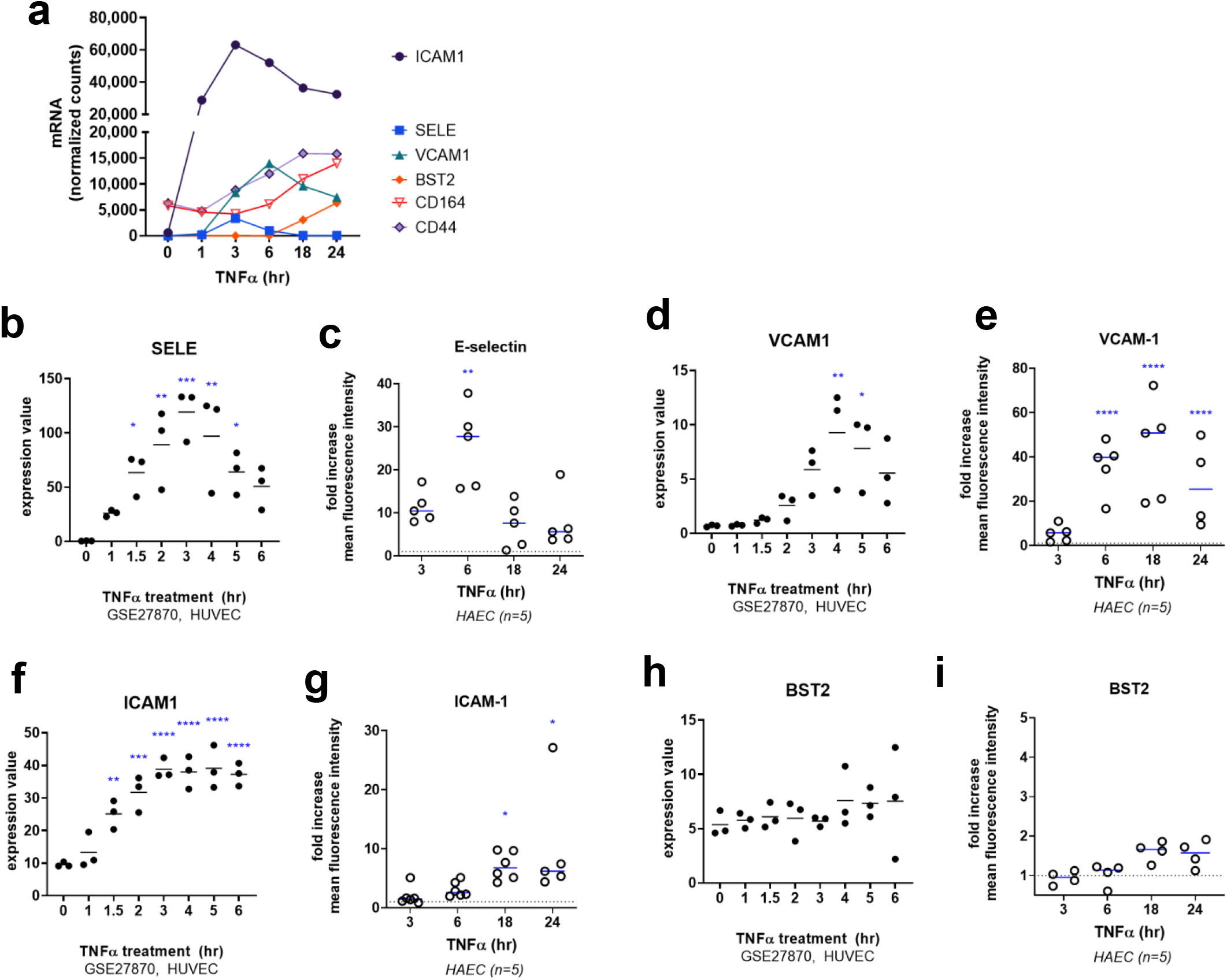
Time course of TNFα stimulated endothelial adhesion molecules. **a**. Time course of TNFα induced adhesion molecule mRNA in HMEC-1 stimulated for 1-24hr. One representative experiment is shown. Results are presented as absolute number of mRNA counts for each gene, normalized to housekeeping controls. **b, d, f, h**. Expression of adhesion molecule mRNA in HUVEC stimulated with TNFα for 1-6hr (GSE27870, n=3). **c, e, g, i** Primary human aortic endothelial cells (HAEC) were stimulated with TNFα (20ng/mL) for 3hr, 6hr, 18hr or 24hr. Cell surface E-selectin (b), VCAM-1 (d), ICAM-1 (f), and BST2 (h) were measured by multiparameter flow cytometry. The fold change in adhesion molecule expression was determined against the MFI of untreated controls (n=4-6 independent experiments). Results are presented as mean fold increase in MFI compared with untreated cells, ± SEM. Expression was compared to untreated endothelium by one way ANOVA followed by Fisher’s LSD. * p<0.05; ** p<0.01; ***p<0.001, **** p<0.0001.

VCAM-1 was also negative in untreated EC. *VCAM1* mRNA was measurable by 3hr of TNFα activation, peaked at 6hr and reduced slightly at 18hr. However, mRNA was maintained at higher levels above baseline through 24hr of TNFα treatment (**Figure 2a, 2d**). VCAM-1 protein was induced by 3hr, peaked at 18hr (72.28±36.65-fold), and was maintained at a slightly lower-than-peak level at 24hr and 40hr, although still elevated compared with untreated conditions (43.6±6.15-fold) (**Figure 2e**).

ICAM-1 mRNA and protein were expressed at a low level in untreated EC. A substantial increase in *ICAM1* transcript was observed within 1hr of TNFα exposure and remained at very high levels through 24hr (**Figure 2a, 2f**). ICAM-1 protein rose modestly at 3hr-6hr and attained its highest levels through 24hr and 40hr of TNFα activation (13.30±6.93-fold) (**Figure 2g**).

*BST2* was lowly detected in untreated EC. mRNA levels did not increase until 18hr after TNFα exposure, but continued to increase at 24hr (**Figure 2a, 2h**). Similarly, BST2 protein was lowly expressed on untreated EC, and increased only later than 6hr of TNFα stimulation. BST2 protein continued to rise through 24hr of TNFα exposure and was maintained at peak levels through 40hr (1.82±0.09-fold) (**Figure 2i**). Other endothelial adhesion molecules *BST1, CD44* and *CD164* (endolyn) were also modestly induced at later time points, with mRNA changes after 6hr and elevated levels maintained at 24hr of TNFα stimulation (**Figure 2a**). In contrast, *ICAM2* was constitutively expressed by EC and was unchanged by activation with TNFα (*data not shown*).

### Time course of TNFα-induced chemokines

TNFα triggered a multi-phase induction of chemokines in endothelium (**Figure 3a**). Chemokines comprising an immediate early response (by 1hr at mRNA level and 3hr at the protein level) in endothelial cells by TNFα were *CXCL1* (GROα), *CXCL2* (MIP-2α), *CXCL3* (MIP-2β) *IL8, CCL2* (MCP-1), *CSF2* (GM-CSF) and *CSF3* (G-CSF). IL-8 and MCP-1 were sustained at high levels through 24hr of stimulation (**Figure 3b, 3c**) and secreted protein accumulated in the supernatant early (**Figure 3d, 3e**). In contrast, endothelial production of early chemokines *CXCL1* (GROα), *CXCL2* (MIP-2α), *CXCL3* (MIP-2β), *CSF2* (GM-CSF) and *CSF3* (G-CSF) declined rapidly over time (**Figure g-n**). A second wave of chemokines was detected later, with comparatively more modest increases not seen until 6hr-18hr: *CX3CL1* (fractalkine), *CCL5* (RANTES), *CCL20* (LARC), *CXCL10* (IP-10), *CXCL11* (I-TAC), and *CSF1* (M-CSF) (**Figure 3p, 3q, 3t, 3u**) with protein increases detected after 6hr of TNFα stimulation (**Figures 3r, 3s, 3v, 3w**). In addition, the tetraspanin *LAMP3*, which is a component of secretory vesicles critical for endothelial intracellular sorting ^5^, was also elevated after 18hr and 24hr of TNFα treatment.

**Figure 3.**
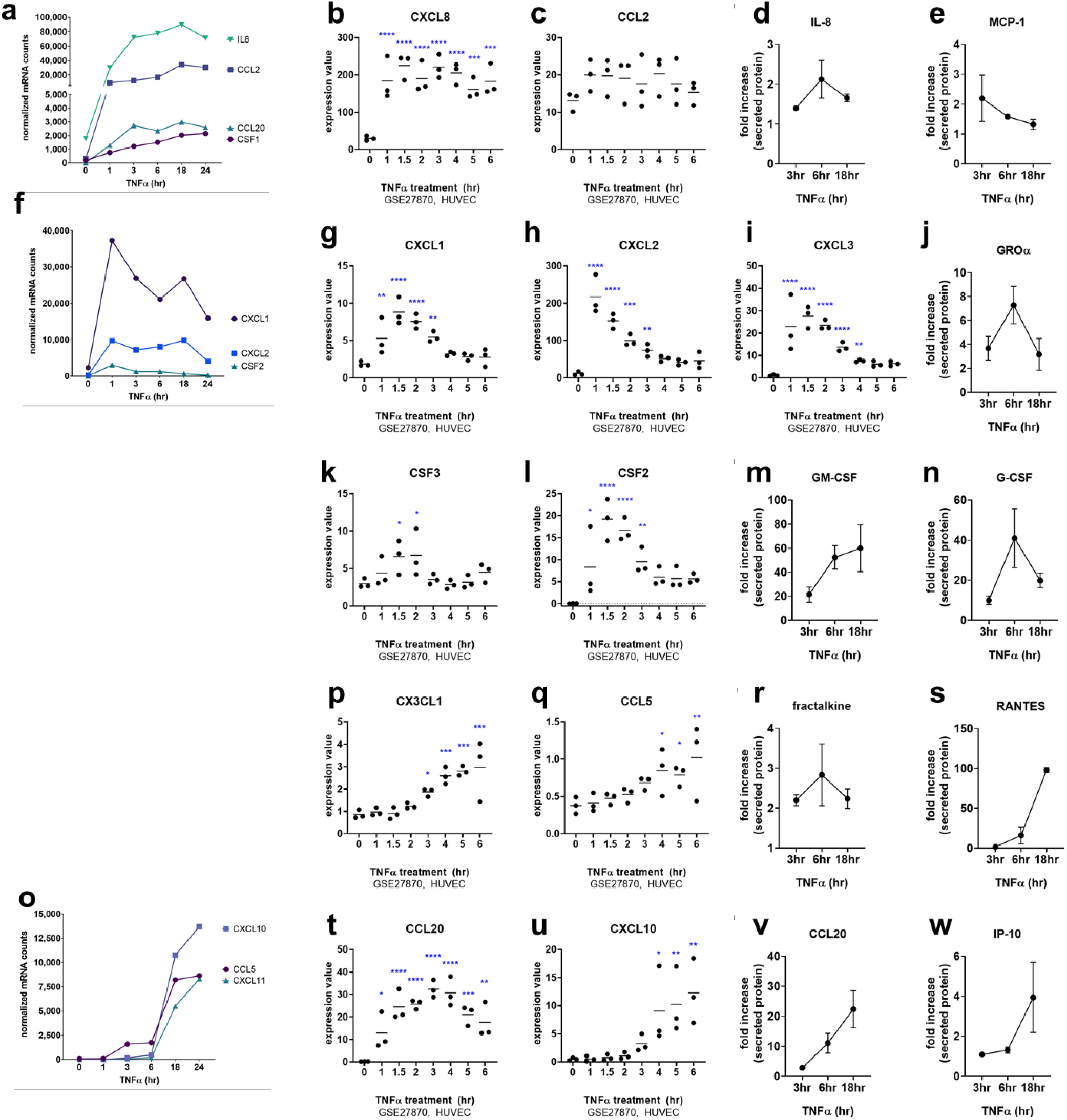
Time course of TNFα stimulated endothelial chemokines. **a-c**. Time course of TNFα induced chemokine mRNA in HMEC-1. One representative experiment is shown. Results are presented as absolute number of mRNA counts for each gene, normalized to housekeeping controls. (a) shows chemokines that were induced early and which declined at later time points; (b) shows chemokines that were induced early and which persisted at high levels for the duration of the experiment; (c) shows chemokines that were induced late. **d-l**. Primary human aortic endothelial cells (HAEC) were stimulated with TNFα (20ng/mL) for 3hr, 6hr, or 18hr. Secreted chemokine protein was measured in the conditioned supernatants by Luminex (panels d-k) and ELISA (CCL20, panel l). Results are presented as mean fold increase in concentration compared to matched untreated controls (n=2 independent experiments). Expression was compared to untreated endothelium by one way ANOVA followed by Fisher’s LSD. * p<0.05; ** p<0.01; ***p<0.001, **** p<0.0001.

### TNFα Withdrawal

To understand the timing of endothelial return to quiescence, we tested cells after different periods of cytokine challenge and extended time of withdrawal from cytokine exposure. Endothelial cells were stimulated with TNFα (20ng/mL) or IFNγ (200U/mL). Then, medium was removed, cells were washed once with M199+10% FBS, and cultured for the remainder of the experiment with fresh M199+10% FBS without cytokine. Adhesion molecules were measured by flow cytometry (**Figure 4a**).

**Figure 4.**
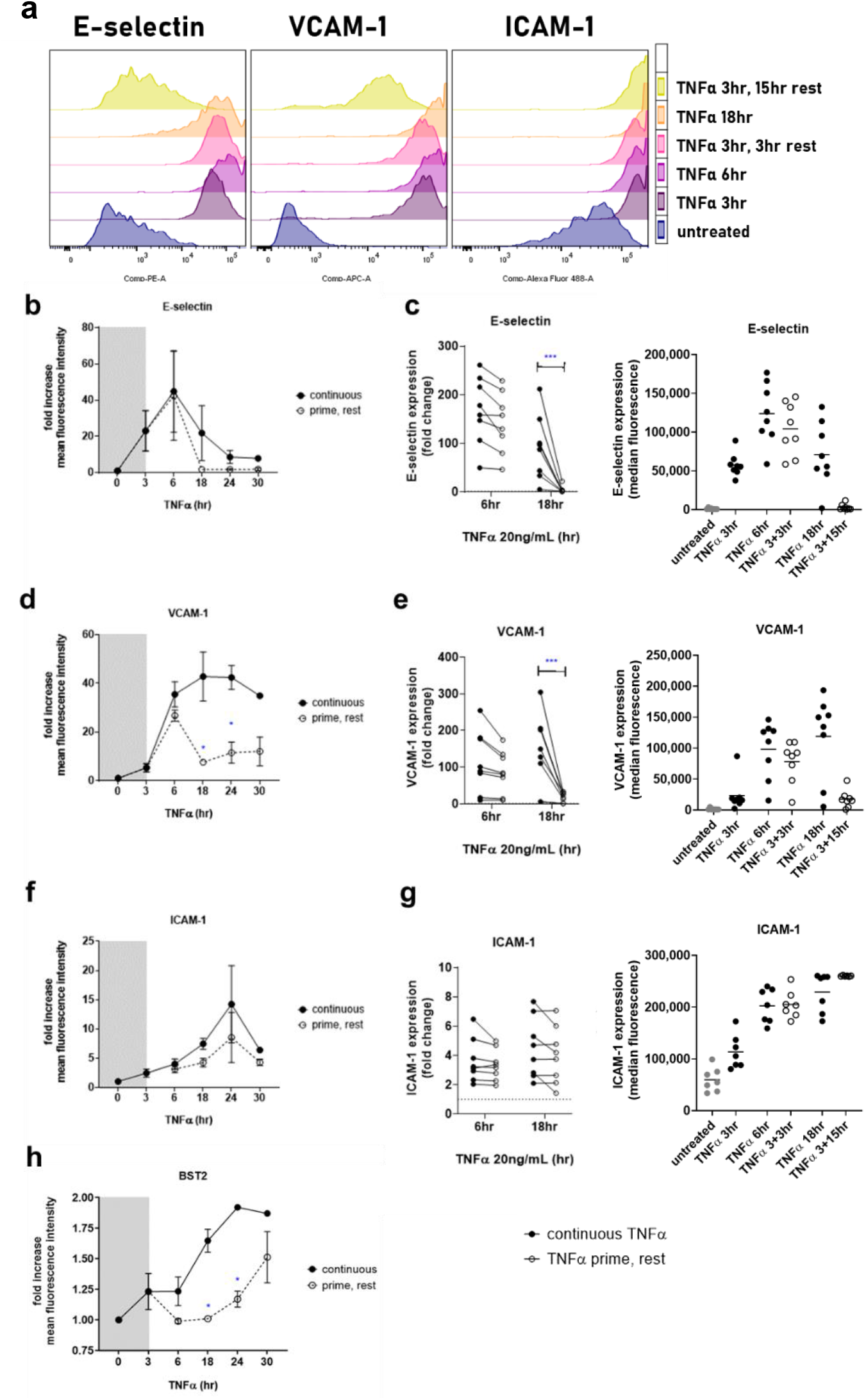
Contracted expression of TNFα-induced adhesion molecules after cytokine withdrawal. Primary endothelia were stimulated with TNFα for 3hr-40hr; or stimulated with TNFα for 3hr and rested in normal medium for an additional 3hr-21hr. Cell surface expression of E-selectin, VCAM-1, ICAM-1 and BST2 was measured by flow cytometry at each time point. **a**. Representative raw flow cytometry data from one experiment with HAEC are shown. **b, d, f, h**. Fold increase in MFI of each adhesion molecule expressed by HAEC, comparing the continuous TNFα presence (closed circles, bold line) to the parallel prime/rest condition (open circles, dashed line). n=5-7 independent experiments. Expression under the continuous vs. prime/rest condition for individual time points were compared by t test, * p<0.05; all other comparisons not significant. **c, e, g**. Primary endothelial cells from 6 different vascular beds were assessed for persistence of adhesion molecule expression. Fold increase is shown in the left panel and raw MFI in the right panel for each marker. Closed circles indicate the fold increase in each adhesion molecule after 6hr or 18hr of TNFα; the open circles indicate the level of adhesion molecule in cells treated for 3hr and then rested for either an additional 3hr or an additional 3hr or 15hr. Paired cells/conditions are graphed in the spaghetti plot. Individual time points were compared by t test, * p<0.05; *** p<0.001; all other comparisons not significant.

We first assessed how long after TNFα stimulation adhesion molecules were maintained at the cell surface. In preliminary experiments, endothelial monolayers were stimulated for 6hr or 18hr with TNFα, then 4hr, 8hr and 24hr after TNFα removal, ICAM-1, VCAM-1 and BST2 were measured. Elevated adhesion molecules were maintained at the same level as at the end of the 18hr stimulation (**Supplemental Figure 2a-2d**), leading us to evaluate shorter priming.

When HAEC were challenged for only 3hr with TNFα, E-selectin protein rose to its peak at 6hr post-exposure and returned to baseline within 15 hours. The kinetics and magnitude of E-selectin induction were nearly identical on HAEC whether TNFα was continuously present over a 30hr period, or was withdrawn after 3hr (**Figure 4b**). Similar results were obtained using endothelial cells from 6 other primary vascular beds (**Figure 4c**). When EC were primed with TNFα for 3hr, E-selection expression peaked at 6hr post-exposure. When allowed to rest for an additional 15hr, E-selectin significantly declined compared with the parallel continuous condition.

VCAM-1 induction followed a similar early pattern, continuing to rise at 6hr post-TNFα exposure. However, VCAM-1 protein decayed by 15hr after TNFα removal. Twenty-four hours after withdrawal, HAEC cell surface VCAM-1 was higher than baseline, but significantly lower than in the presence of continuous TNFα (reaching 27.1±7.3% of maximum at 24hr) (**Figure 4d**). Similar patterns were observed with 6 other vascular beds (**Figure 4e**), where VCAM-1 expression returned to just above baseline at 18hr, after 3hr priming with TNFα.

In contrast, ICAM-1 protein expression reached near peak levels on HAEC regardless of whether TNFα was present for 30hr, or cells were only primed for 3hr (ICAM-1, 63.5±14.3% of max at 24hr) (**Figure 4f**). Elevated ICAM-1 expression was also maintained on the cell surface of 6 other endothelial cell types, and was not significantly different from the same time point in the continuous presence of TNFα (**Figure 4g**). Like ICAM-1, BST2 protein expression was significantly elevated at later time points after only 3hr TNFα exposure of TNFα (BST2, 70.9±9.6% of max at 24hr) (**Figure 4h**).

We also measured mRNA levels of adhesion molecules after 3hr priming with TNFα, followed by withdrawal for an additional 3hr, 15hr or 21hr, and compared them to continuous TNFα exposure at the same time points (**Supplemental Figure 3**). The kinetics of *SELE* (E-selectin) mRNA were identical in both conditions (**Supplemental Figure 3a**), whereas *VCAM1* mRNA plateaued 3hr after withdrawal and then rapidly returned to baseline (**Supplemental Figure 3b**). *ICAM1* mRNA contracted rapidly by 18hr after TNFα exposure, unlike the pattern for protein expression which was persistently high through 40hr (**Supplemental Figure 3c**); it is possible that ICAM-1 protein is highly stable or minimally recycled compared with other adhesion molecules. In contrast, *BST2* and *CD164* mRNA did increase at 18hr after only 3hr priming, although the levels were lower than at the same time point when TNFα was continuously present (**Supplemental Figures 3d-3f**).

Unlike ICAM-1 and BST2, TNFα-induced chemokines required a continuous presence of TNFα to be produced at maximal amounts. Some chemokines, *IL8, CCL2* (MCP-1), *CCL20* (LARC), *CSF2* (GM-CSF), declined to negligible levels when TNFα was no longer present. Others were persistently produced at low levels up to 24hr after TNFα had been removed, including *CCL5* (RANTES), *CXCL1* (GROα), *CXCL2* (MIP-2α), and *CXCL10* (IP-10) (**Figure 5;** mRNA counts in **Supplemental Figure 4**). But, in all cases, chemokine secretion was significantly lower from EC with only short exposure to TNFα compared with continuous stimulation.

**Figure 5.**
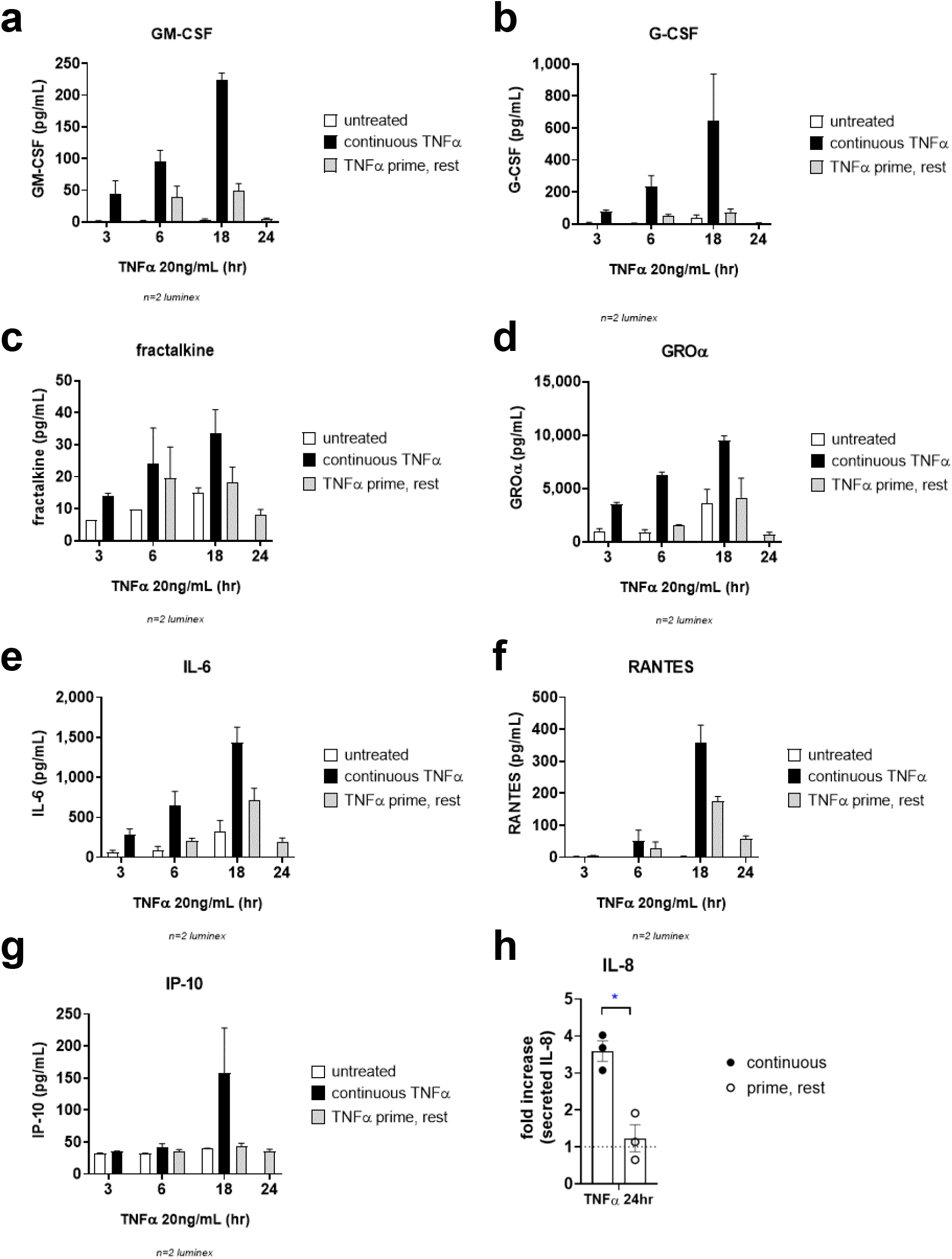
Rapidly declining secretion of TNFα-induced chemokines after cytokine withdrawal. **a-h**. HAEC were stimulated with TNFα for 3hr, 6hr, 17hr or 24hr (continuous, black bars); or primed with TNFα for 3hr, followed by cytokine withdrawal for 3hr, 15hr or 21hr (prime/rest, grey bars). Conditioned medium from untreated cells was also collected at each time point (white bars). Supernatants were assayed for secreted chemokines by Luminex (panels a-g, n=2) and ELISA (IL-8, panel h, n=3). Data are presented as fold increase in chemokine concentration (pg/mL) normalized to untreated, mean+/- SEM. Individual time points were compared by t test, * p<0.05.

In summary, as little as 3hr of TNFα exposure triggered long-term changes in endothelial cell phenotype, with elevated ICAM-1 and BST2 expression 24hr later. However, the repertoire was less broad after TNFα withdrawal compared with chronic TNFα stimulation, with lower VCAM-1 and little E-selectin or chemokine production, and low level sustained expression of other pro-adhesive factors.

### IFNγ-induced adhesion molecules

We next examined the kinetics and durability of endothelial response to IFNγ. We measured mRNA in HMEC-1 treated with IFNγ for 1, 3, 6, 18 and 24hr (**Figure 6a**); and also examined a public dataset of primary human lung endothelium (HLVMEC) treated with IFNγ for 3, 6 and 24hr (GSE106524). We further confirmed adhesion molecule and chemokine expression in primary human aortic endothelium (n=4) and primary human endothelial cells from 6 other vascular beds (coronary artery, cardiac microvasculature, pulmonary artery, lung microvasculature, renal microvasculature, dermal microvasculature) (**Figure 6b**).

**Figure 6.**
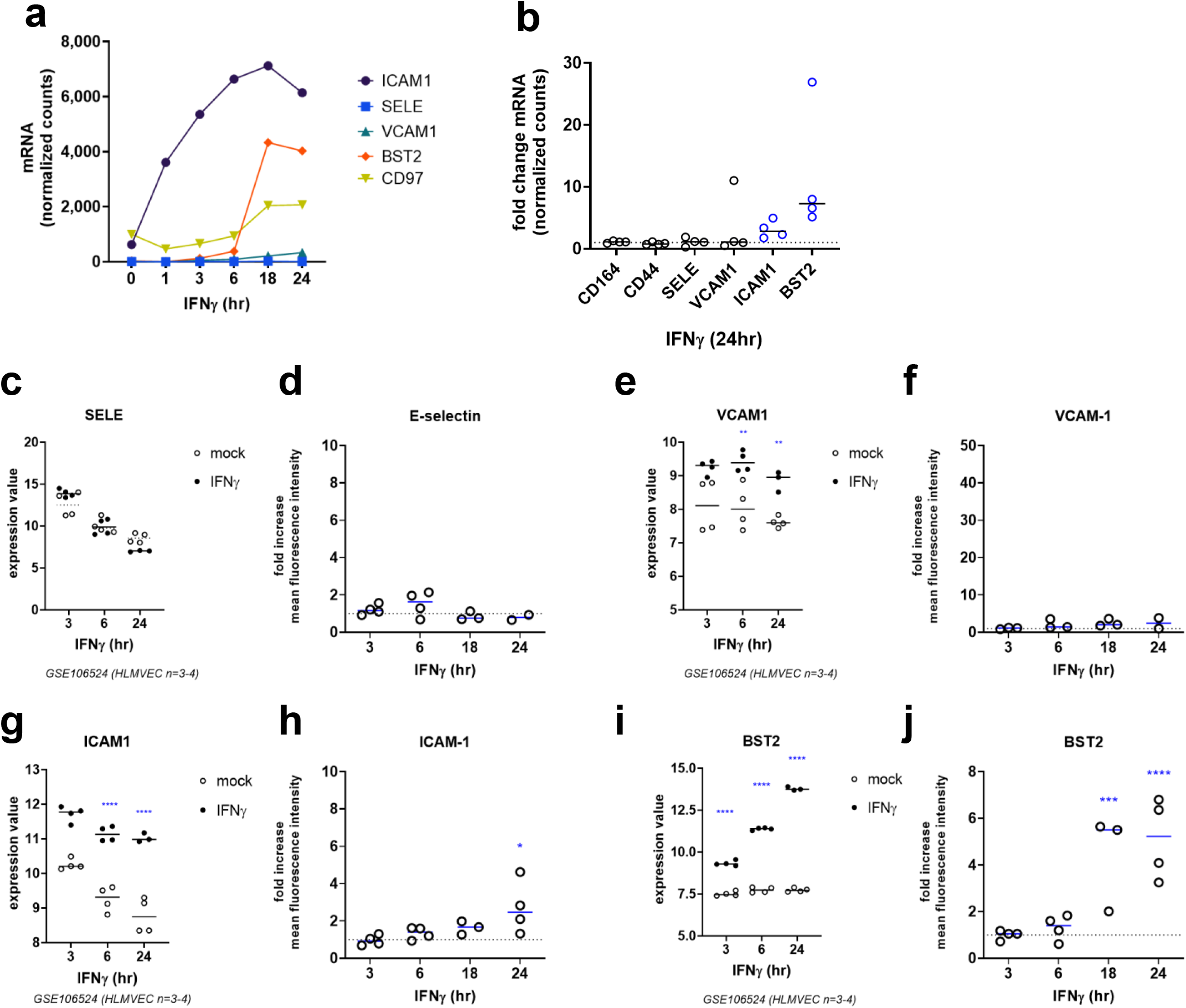
Time course of IFNγ stimulated endothelial adhesion molecules. **a**. Time course of IFNγ induced adhesion molecule mRNA in HMEC-1. One representative experiment is shown. Results are presented as absolute number of mRNA counts for each gene, normalized to housekeeping controls. **b**. Fold change in adhesion molecule transcripts induced by IFNγ (24hr) across 4 different primary endothelial cells. Results are presented as fold increase compared to each cell’s untreated parallel control. Blue symbols indicate mean increase ≥2-fold. **c, e, g, i**. Time course of adhesion molecule mRNA in primary human lung microvascular endothelium stimulated with IFNγ for 3hr, 6hr or 24hr (n=4, GSE106524). Expression values for *SELE* (c), *VCAM1* (e), *ICAM1* (g), and *BST2* (i) are shown. IFNγ (closed circles) was compared to the mock parallel control (open circles) at each time point. **d, f, h, j**. Primary human aortic endothelial cells were treated with IFNγ (200U/mL) for 3hr, 6hr, 18hr or 24hr. Cell surface E-selectin (g), VCAM-1 (h), ICAM-1 (i), and BST2 (j) were measured by multiparameter flow cytometry. Results are presented as mean fold increase in MFI compared with untreated cells, ±SEM (n=3-4 independent experiments). Two way ANOVA, vs. 3hr ** p<0.01; **** p<0.0001 compared to paired untreated controls.

Unlike TNFα stimulation, E-selectin was not induced on any EC (**Figure 6a, 6b, 6c**). VCAM-1 expression by IFNγ was modest and variable across experiments and endothelial cell types (**Figure 6a, 6b, 6e, 6f**). ICAM-1 was slightly but consistently increased by IFNγ, starting at 18hr, rising to 2.71±0.6-fold at 24hr on HAEC and 3.09±0.7-fold across other EC (**Figure 6a, 6b, 6g, 6h**). IFNγ most robustly stimulated BST2 expression, with increased mRNA and protein detected at 18hr, rising to 5.13±0.8-fold at 24hr on HAEC (n=4) (**Figure 6a, 6b, 6j**). Similarly, BST2 was induced at 24hr at the mRNA level by 11.65±5.1-fold (**Figure 6b**) and protein level at 1.93±0.27-fold (**Figure 6j**) across five other EC types.

The EC adhesion molecule *CD97* ^6^ was slightly increased at the transcript level by IFNγ at 18hr and 24hr, but IFNγ had no effect on *CD164, CD44, BST1* or *ICAM2* (data not shown).

### IFNγ-induced chemokines

The range of IFNγ-induced endothelial chemokines was narrower and mostly distinct from that induced by TNFα. Predominant signatures were *CXCL9* (MIG), *CXCL10* (IP-10), and *CXCL11* (I-TAC), mRNA for which emerged by 3hr and peaked at 18hr (**Figure 7a**). *CXCL9* (MIG), *CXCL10* (IP-10), and *CXCL11* (I-TAC) transcripts were prominently induced across multiple primary endothelial cell types (Figure **7b, 7c, 7d, 7e**). IP-10 and I-TAC protein was detected in conditioned supernatants of HAEC as early as 6hr and rose beyond 24hr (**Figure 7f, 7g**).

**Figure 7.**
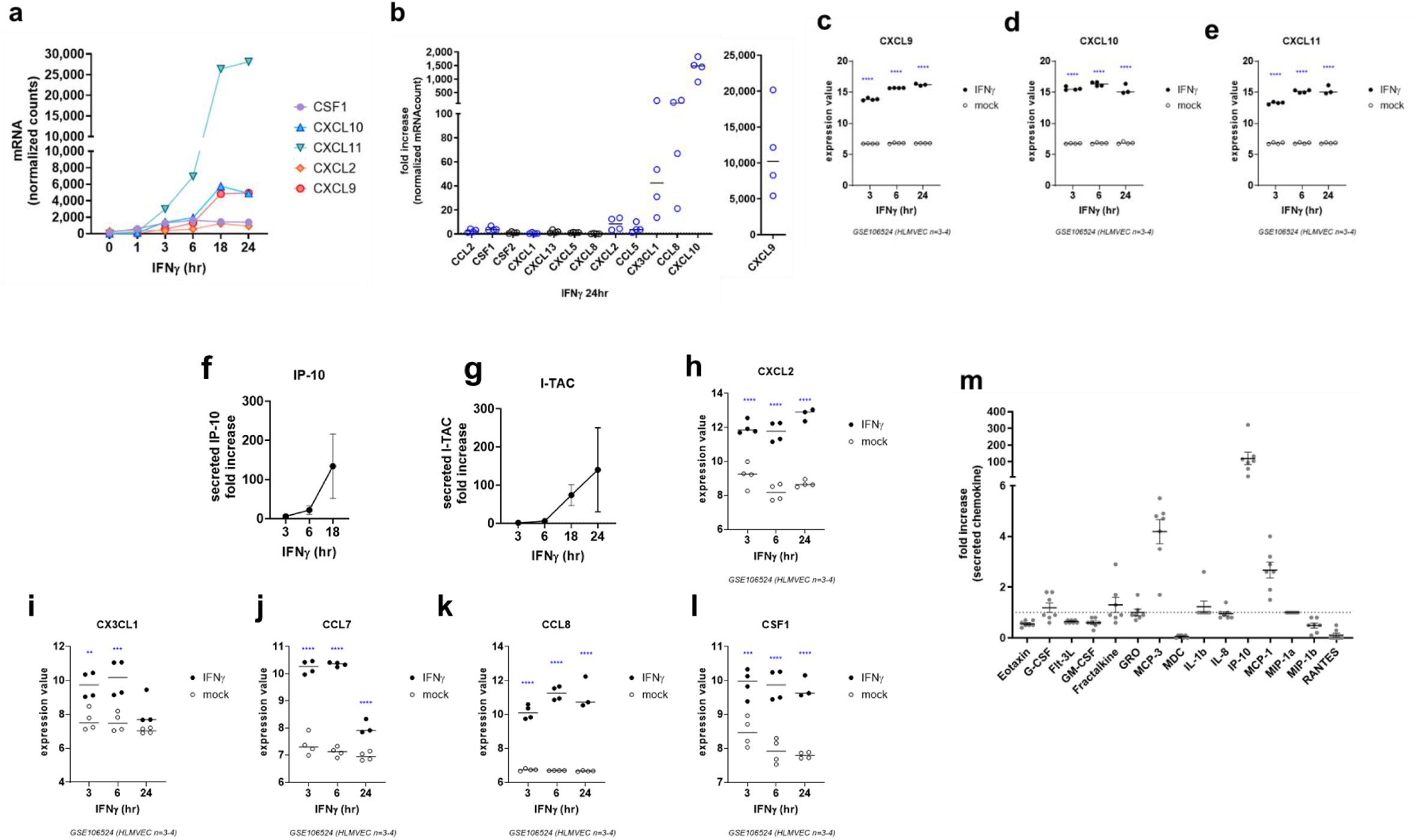
Time course of IFNγ stimulated endothelial chemokines. **a**. Time course of IFNγ induced chemokine mRNA in HMEC-1. One representative experiment is shown. Results are presented as absolute number of mRNA counts for each gene, normalized to housekeeping controls. **b**. Fold change in chemokine transcripts induced by IFNγ (24hr) across 4 different primary endothelial cells. Results are presented as fold increase compared to each cell’s untreated parallel control. Blue symbols indicate mean increase ≥2-fold. **c-e**. Time course of *CXCL9* (c), *CXCL10* (d), and *CXCL11* (e) chemokine mRNA in primary human lung microvascular endothelium stimulated with IFNγ for 3hr, 6hr or 24hr (n=4, GSE106524). Expression values are shown. IFNγ (closed circles) was compared to the mock parallel control (open circles) at each time point. Two way ANOVA, **** p<0.0001 compared to paired untreated controls. **f, g**. Primary human aortic endothelial cells were treated with IFNγ for 3hr, 6hr, 18hr or 24hr. Secreted chemokine protein was measured in the conditioned supernatants by Luminex (IP-10) and ELISA (I-TAC). Results are presented as mean fold increase in secreted IP-10 (f) and I-TAC (g) chemokines from IFNγ-stimulated HAEC (n=3 independent experiments), mean ±SEM. **h-l**. GSE106524, time course of chemokine mRNA in primary human lung microvascular endothelium stimulated with IFNγ for 3hr, 6hr or 24hr (n=4, GSE106524). Expression values for each gene are shown. IFNγ (closed circles) was compared to the mock parallel control (open circles) at each time point. Two way ANOVA, **** p<0.0001 compared to paired untreated controls. **m**. Primary human endothelial cells from 7 vascular beds were treated with IFNγ for 24hr. Secreted chemokine protein was measured in the conditioned supernatants by Luminex. Results are presented as fold increase in secreted chemokine in medium, showing one representative experiment of three.

In addition, IFNγ consistently induced low levels of *CCL2* (MCP-1), *CXCL1* (GROα), MCP-3 (*CCL7*), *CXCL2* (MIP-2α), *CCL5* (RANTES), *CX3CL1* (fractalkine), and *CCL8* (MCP-2) mRNA in endothelial cells from four primary vascular beds (**Figure 7b, 7h, 7i, 7j, 7k, 7l**, discrete mRNA counts across each cell type are provided in **Supplemental Figure 5**). Increased chemokine secretion of G-CSF (*CSF3*), fractalkine (*CX3CL1*), MCP-3 (*CCL7*), and MCP-1 (*CCL2*) was confirmed from primary endothelial cells after IFNγ stimulation for 24hr (**Figure 7m**). There were no changes in other CCL and CXCL chemokines or *IL8* caused by IFNγ stimulation (**Figure 1a**).

### IFNγ Withdrawal

We next assessed short-term exposure to IFNγ, where endothelium was primed with IFNγ for 3hr, then normal medium was added for the remainder of the experiment. Representative flow cytometry data of cell surface adhesion molecule expression are shown in **Figure 8a**.

**Figure 8.**
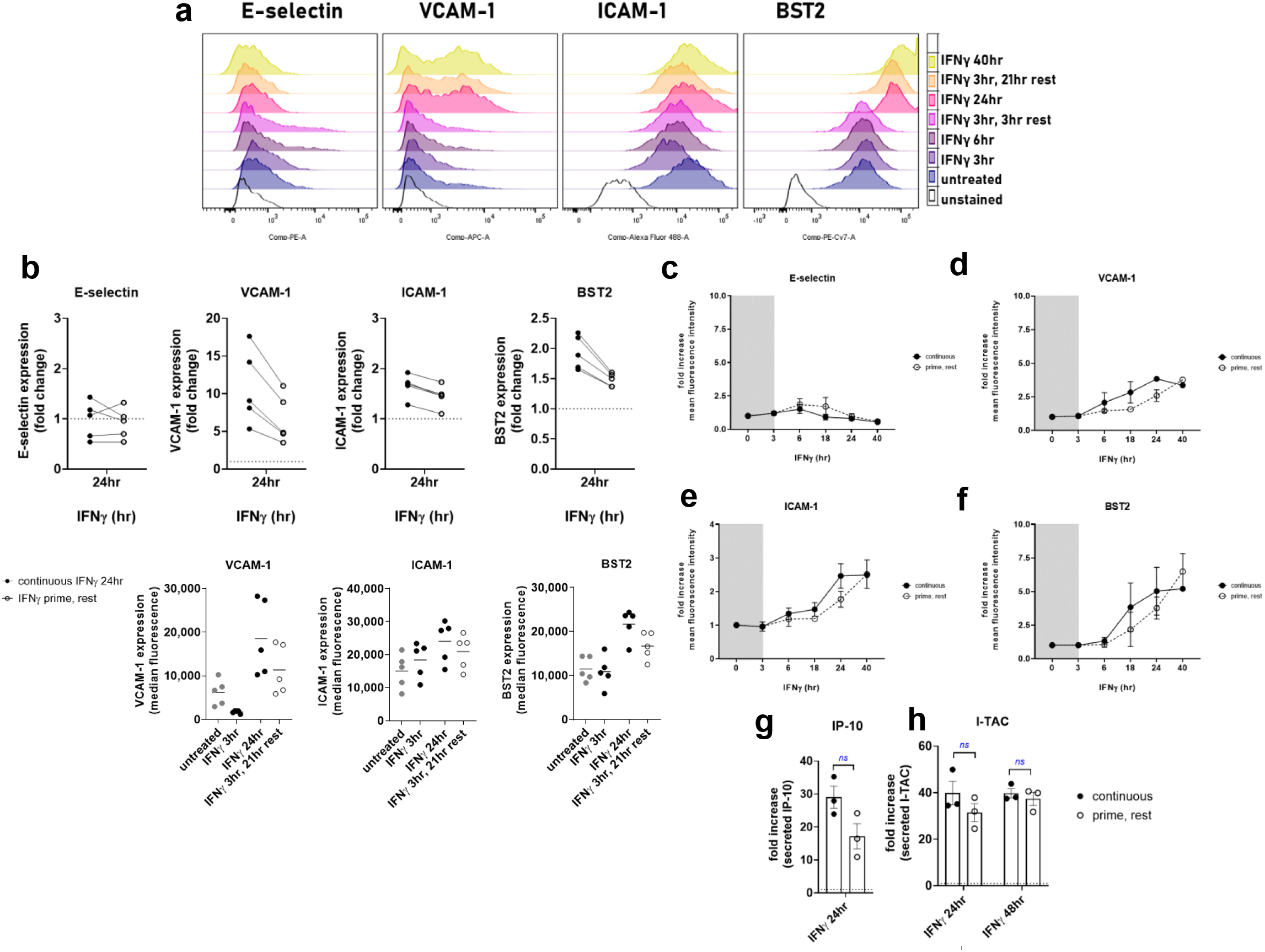
Persistent expression of IFNγ-induced adhesion molecules and chemokines after cytokine withdrawal. Primary human endothelial cells were stimulated with IFNγ for 3hr, 6hr, 18hr, 24hr or 40hr; or stimulated with IFNγ for 3hr and rested in normal medium for an additional 3hr, 15hr, or 21hr. Cell surface expression of E-selectin, VCAM-1, ICAM-1 and BST2 was measured by flow cytometry at each time point. Secreted chemokines were measured in the supernatant by ELISA. **a**. Representative raw flow cytometry data from HAEC are shown. **b**. Primary endothelial cells from 5 different vascular beds were assessed for persistence of adhesion molecule expression. Closed circles indicate the fold increase in each adhesion molecule after 24hr of continuous IFNγ; the open circles indicate the level of adhesion molecule in cells treated for 3hr and then rested for 21hr. Paired cells/conditions are graphed in the spaghetti plot. **c-f**. Fold increase in the MFI of each adhesion molecule expressed by HAEC, comparing the continuous IFNγ presence (closed circles, bold line) to the parallel prime/rest condition (open circles, dashed line), n=3-4 independent experiments. Results are expressed as the fold increase over untreated, paired cells/conditions are graphed in the spaghetti plot (n=5). Expression under the continuous vs. prime/rest condition for individual time points were compared by t test, all comparisons not significant **g, h**. Persistence of secreted chemokines IP-10 (**g**) and I-TAC (**h**) after 3hr IFNγ priming and withdrawal for an additional 21hr. n=3 HAEC. Expression under the continuous vs. prime/rest condition for individual time points were compared by t test, all comparisons not significant

As above, continuous IFNγ did not induce cell surface E-selectin (**Figure 8b, 8e**) and slightly increased VCAM-1 (**Figure 8c, 8d**). Unlike with TNFα, however, priming of EC with IFNγ for 3hr resulted in elevated levels of VCAM-1, BST2 and ICAM-1 21hr later (**Figure 8b**, n=5 primary EC). In particular, BST2 and ICAM-1 were strongly induced at 18hr, 24hr and 40hr. Moreover, the magnitude of expression after IFNγ was removed was not significantly different from that under continuous presence of IFNγ up to 40hr later for both ICAM-1 (**Figure 8e**) or BST2, which reached 99.45±0.5% of maximum (**Figure 8f**). mRNA counts are shown in **Supplemental Figure 6a-6e**.

Similarly, the amount of *CXCL10* (IP-10) and *CXCL11* (I-TAC) produced by EC was not significantly different after only 3hr exposure to IFNγ compared with continuous IFNγ (fold change is shown in **Figure 8g, 8h**; absolute concentrations provided in **Supplemental Figure 6f, 6g**). The same results were obtained with four other primary endothelial cells, where secretion of IP-10 and I-TAC was substantially elevated even if endothelium were only primed for 3hr with IFNγ (**Figure 8i, 8j**). Even as short as 1hr or 2hr exposure to IFNγ caused maximal IP-10 secretion and BST2 expression 24hr later (**Supplemental Figure 6f**). Other chemokines *CXCL9* (MIG), *CCL7* (MCP-3) and *CCL2* (MCP-1) remained increased over control after IFNγ priming and withdrawal, albeit at lower than peak levels (**Supplemental Figure 6h-6l**).

Hence, in contrast to TNFα, short exposure of endothelial cells to IFNγ was sufficient to induce an enhanced inflammatory phenotype. In particular BST2 and IP-10 exhibited markedly high expression on endothelium up to 48hr after IFNγ was last present.

### Differential Requirements for Intracellular Signaling Pathways in Endothelial Activation

We wondered whether differential intracellular signaling pathways accounted for the long-term durability of IFNγ effects compared with TNFα.

TNFα canonically exerts its effects through NFκB. The time course of NFκB-associated genes is presented in **Figure 9a**, with strong TNFα induction of *NFKBIA* (IκBα), *TNFAIP3* (A20) and *BCL3* by 1hr, and *NFKB1* (p105/p50), *NFKB2* (p100/p52), *RELA* (p65), *RELB*, and *NFKBIZ* (IκBζ) by 3hr. While *BCL3* (an atypical transcription activator) mRNA increases were transient, all other NFKB-related genes remained elevated through 24hr of continuous TNFα exposure (**Figure 9a, 9b**). At the mRNA and protein level, NFκB inhibition with MG132 blocked TNFα induced E-selectin (95.5±3.9%, **Figure 9d, 9f**), VCAM-1 (99.0±0.8%, **Figure 9d, 9g**) and ICAM-1 (84.4±13.7%, **Figure 9d, 9h**). NFκB was also required for TNFα triggered chemokine expression, including IL-6 (**Figure 9l**), MCP-1 (**Figure 9m**), GROα (**Figure 9n**), IL-6 (**Figure 9o**), GM-CSF (**Figure 9p**) and G-CSF (**Figure 9q**), but did not have a significant effect on BST2 (−9.7±9.4%, **Figure 9d, 9i**, raw MFI in **Supplemental Figure 7a-7d**).

**Figure 9.**
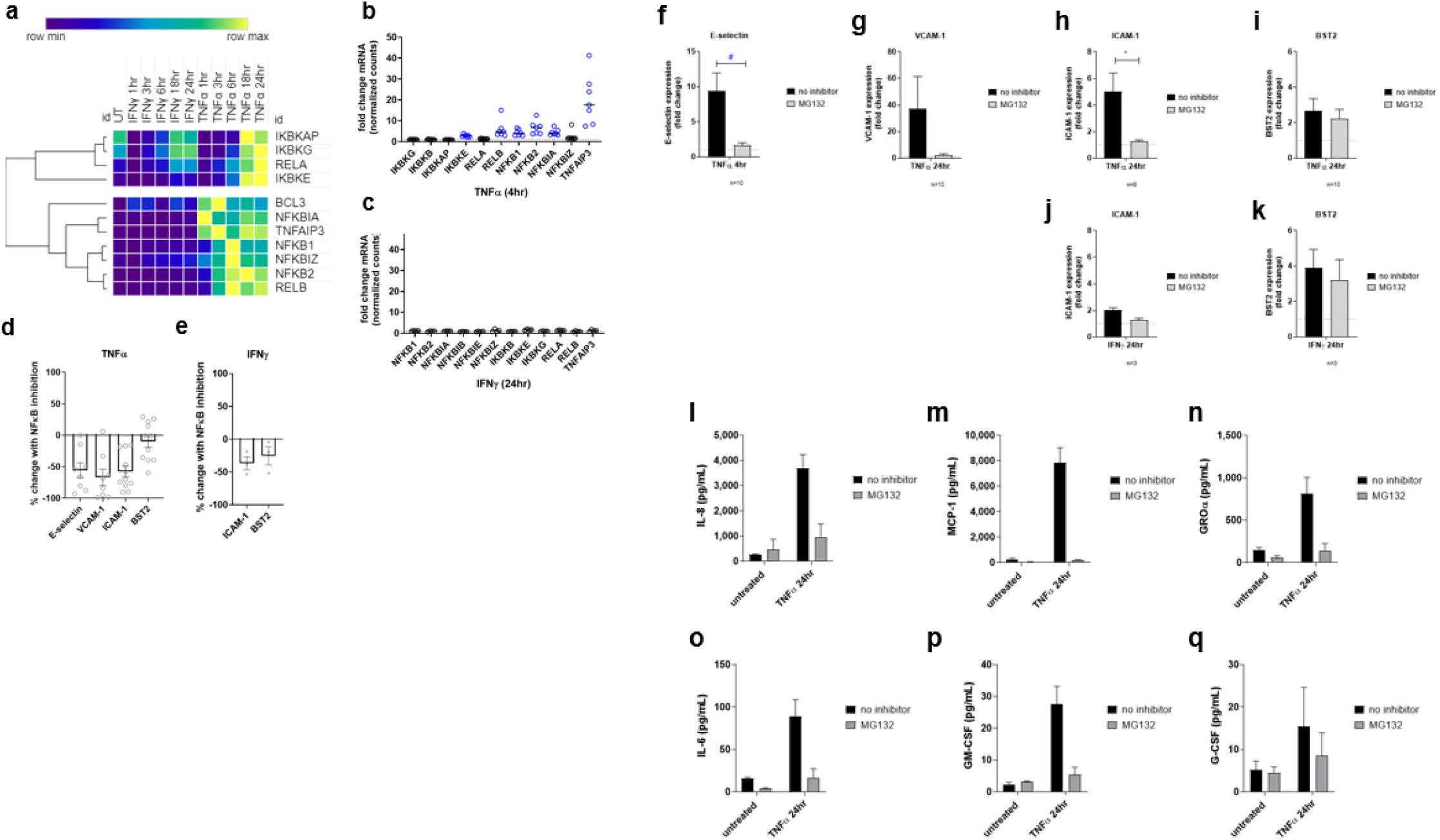
NFκB dependent endothelial cell activation. HAEC were left uninhibited or pre-treated with the NFκB inhibitor MG132 (5μM) for 30min, prior to continuous stimulation with TNFα. Cell surface adhesion molecules were measured by flow cytometry, and secreted chemokines were measured by Luminex and ELISA. **a**. Heat map shows relative transcript counts of NFκB related genes in HMEC-1 stimulated with IFNγ or TNFα for 1-24hr, measured by Nanostring. **b, c**. Fold change in NFκB-related transcripts induced by TNFα (4hr, panel b) or IFNγ (24hr, panel c) across 6 different primary endothelial cells. Results are presented as fold increase compared to each cell’s untreated parallel control. Blue symbols indicate mean increase ≥2-fold. **d, e**. HAEC were pre-treated with MG132 (5μM) for 30min, prior to stimulation with TNFα (panel d, n=8 independent experiments) or IFNγ (panel e, n=3). Cell surface adhesion molecule expression was measured by flow cytometry. Graph shows mean percent inhibition by NFκB inhibitor MG132 of E-selectin (at 4hr), VCAM-1, ICAM-1 and BST2 (at 24hr) protein induction. **f-i**. TNFα-induced cell surface E-selectin (f), VCAM-1 (g), ICAM-1 (h), and BST2 (i) expression without inhibitor (black bars) and with MG132 (grey bars). Results are shown as mean fold change in MFI ±SEM, n=9-10 independent experiments. Conditions were compared by t test, #p<0.1, * p<0.05. **j-k**. Fold change in IFNγ-induced cell surface expression of ICAM-1 (j) and BST2 (k) without inhibitor (black bars) and with MG132 (grey bars). Results are shown as mean fold change in MFI ±SEM, n=3 independent experiments. Conditions were compared by t test, neither comparison significant. **l-q**. Fold change in TNFα-induced chemokine secretion without inhibitor (black bars) and with MG132 (grey bars). Results show concentration of each chemokine in conditioned medium, mean ±SEM (n=2 HAEC).

In contrast, no NFκB associated genes were altered >2-fold by IFNγ stimulation (**Figure 9a, 9c**). Unsurprisingly, NFκB inhibition did not affect IFNγ induction of ICAM-1 or BST2 protein expression (**Figure 9e, 9j, 9k**).

TNFα is known to trigger phosphorylation and activation mitogen-activated protein kinase (MAPK) pathways, including Jun N-terminal kinase (JNK), p38 and ERK ^7,8^. We also observed regulation of MAPKs at the transcript level in EC by TNFα. Continuous TNFα treatment of HMEC-1 elevated mRNA expression of *MAP4K4* (upstream JNK activator), *MAP4K2* (upstream JNK activator), *MAPK1* (ERK2), *MAPK11* (p38β), and *MAPKAP2* (downstream p38 target), but not *MAP4K1* (HPK1), all beginning at 6hr and sustained to 24hr (**Supplemental Figure 8a**). MAPK gene expression remained high when cells were primed for 3hr with TNFα and rested in the absence of cytokine, up to 18hr, with transcript levels beginning to decline at 24hr (**Supplemental Figure 8b**). In response to IFNγ, however, only *MAPK1* (ERK2) was elevated (**Supplemental Figure 8c, 8d**). We next pre-treated primary endothelium with the JNK inhibitor (SP600125, 20μM), p38 inhibitor (SB203580, 25μM) or MEK/ERK inhibitor (U0126, 10μM) prior to stimulation with TNFα and measurement of adhesion molecules and chemokine expression. p38 but not JNK or ERK was required for E-selectin (p38: 64.76±12.0% inhibition). VCAM-1 induction was also primarily p38-dependent (p38: 83.5±7.0% inhibition), while TNFα-induced BST2 and ICAM-1 were not significantly affected by any MAPK inhibitor (**Supplemental Figure 8e**). Among TNFα-induced chemokines, secretion of IL-6, IL-8, MCP-1, RANTES was blocked by the p38 inhibitor; IP-10 was suppressed by the JNK but not p38 inhibitor; and both p38 and JNK were required for maximal GM-CSF, G-CSF, and fractalkine (**Supplemental Figure 8f**).

We next examined JAK/STAT and interferon response gene expression under TNFα and IFNγ stimulation. TNFα strongly increased expression of *IRF1* and showed modest increases in *STAT1, STAT5A* and *IRF7*, but did not significantly affect other interferon response genes (**Figure 10a**). We confirmed early and robust *IRF1* expression in primary endothelial cells at 4hr from 6 vascular beds (**Figure 10b**) and a public dataset of HUVEC stimulated with TNFα for 1-6hr (GSE27870), where *IRF1* peaked between 1.5hr and 3hr, and tapered off by 6hr (**Figure 10c**). While *STAT1* was only slightly changed (**Figures 10b, 10d**), *STAT5A* transcript increased early in both of our datasets (**Figure 10a, 10b**) and in GSE27870 (**Figure 10e**), and remained elevated through 24hr.

**Figure 10.**
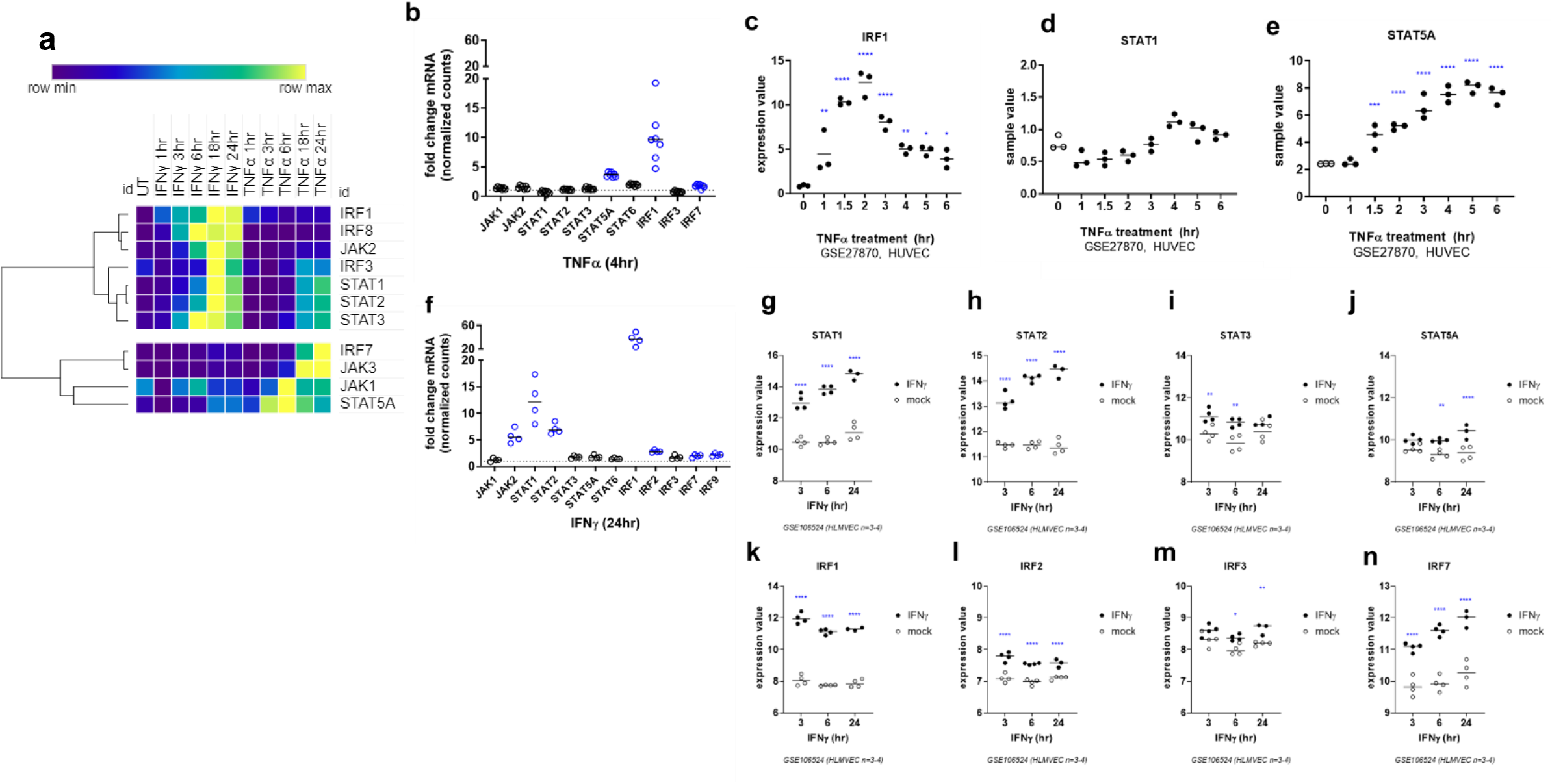
Differential induction of JAK/STAT signaling pathways by TNFα and IFNγ in endothelial cells. **a**.Heat map shows relative transcript counts of JAK, STAT and IRF genes in HMEC-1 stimulated with IFNγ or TNFα for 1-24hr, measured by Nanostring. **b**.Fold change in JAK, STAT and IRF transcripts induced by TNFα (4hr) across 6 different primary endothelial cells. Results are presented as fold increase compared to each cell’s untreated parallel control. Blue symbols indicate mean increase ≥2-fold. **c-e**. HUVEC were treated with TNFα for 1-6hr (n=3, GSE27870). Transcript expression values of *IRF1* (c), *STAT1* (d), and *STAT5A* (e) are shown. Expression at each time point was compared to the untreated condition by one way ANOVA. **f**. Fold change in JAK, STAT and IRF transcripts induced by IFNγ (24hr) across 4 different primary endothelial cells. Results are presented as fold increase compared to each cell’s untreated parallel control. Blue symbols indicate mean increase ≥2-fold. **g-n**. HLMVEC were treated with IFNγ for 3hr, 6hr or 24hr (n=4, GSE106524). The time course of STAT and IRF mRNA in primary human lung microvascular endothelium is shown. IFNγ (closed circles) was compared to the mock parallel control (open circles) at each time point. Expression at each time point was compared to the untreated condition by one way ANOVA. * p<0.05; ** p<0.01; ***p<0.001; **** p<0.0001.

IFNγ robustly augmented *JAK1, STAT1*, and *STAT2* gene expression in HMEC-1 (**Figure 10a**) and primary endothelial cells (**Figure 10f**). IFNγ additionally induced *JAK2* and other interferon response factors, most highly *IRF1*. The timing of STAT and IRF transcript changes was confirmed in a public dataset of primary lung endothelial cells (GSE106524), where IFNγ treatment prominently elevated *STAT1* and *STAT2*, but not *STAT3* or *STAT5A* (**Figures 10g-j**); and *IRF1, IRF2* and *IRF7*, but not *IRF3* (**Figures 10k-n**).

Therefore, at the transcriptional level, TNFα triggered a predominant NFκB signature cooperatively supported by p38 MAPK, while IFNγ drove broad JAK/STAT programming. However, *STAT* genes and *IRF1* were common to both TNFα and IFNγ in endothelial cells.

In IFNγ signaling, STAT1 homodimers bind to GAS motifs, within regulatory regions of genes such as SOCS3 and IRF1. Induction of IRF1 constitutes a second wave of transcription activation, binding to genes harboring IRSE/IRE motifs. To evaluate binding of STATs, a ChIP-seq dataset in HeLa and K562 cells treated with IFNγ for 30min or 6hr was analyzed from ENCODE. We also determined the presence of transcription factor motifs in the *SELE, VCAM1, ICAM1, BST2*, and *CXCL10* genes using MotifMap ^9,10^ and analyzed ChIP-Seq data for NFκB, STAT and IRF binding to these genes (**Supplemental Table 1**). The *ICAM1* gene is not only NFκB responsive but also contains a GAS motif, putatively regulated by STAT1; and is bound by IRF1 (per K562 ENCODE) (in addition to NFκB/RELA). Similarly, *CXCL10* contains both ISRE and NFκB binding sites. *BST2* has a composite GAS and ISRE motif that could be regulated by both STAT1 and IRF1 (especially the STAT1:STAT2:IRF9 complex ISGF3) ^11^, but no NFκB motifs. *SELE* and *VCAM1* are likely regulated only by NFκB, but not STAT1 or IRF1.

We therefore dissected the functional requirement for JAK in endothelial cell activation (**Table 1**). Endothelial cells were pre-treated with JAK1/2 inhibitor ruxolitinib, before stimulation with TNFα or IFNγ. At the protein level, JAK inhibition had no effect on TNFα induced E-selectin (**Figure 11a, 11c**), ICAM-1 (**Figure 11a, 11d**), or VCAM-1 (**Figure 11a, 11f**), but partially reduced BST2 (45.5±14.9%, **Figure 11a, 11e**). Raw MFI results are given in **Supplemental Figure 9a-9f**.

**Table 1.** Regulation of selected genes by STAT, IRF and NFκB

| Gene | GAS Motif (STAT) | ISRE/IRE Motif (IRF) | NFκB Motif |
| --- | --- | --- | --- |
| *SELE* | n | IRF8 | y |
| *VCAM1* | n | n | y |
| *ICAM1* | y* | n | y |
| *BST2* | y (ISGF3) | y (IRF1, IRF2, IRF8) | n |
| *CXCL10* | y (ISGF3) | y (IRF1, IRF2, IRF8) | y |

**Table 2.** Percent inhibition of inducible adhesion molecule expression by JAK1/2 inhibitor ruxolitinib. Results are mean ± SEM.

| Protein | TNFα | IFNγ |
| --- | --- | --- |
| ICAM-1 | -10.3±12.67% | -53.8±6.7% |
| E-selectin | 3.28±26.47% | [-3.517±14.9]% |
| VCAM-1 | -7.22±14.5% | -63.13±14.75% |
| BST2 | -45.46±14.9% | -68.49±6.69% |

**Figure 11.**
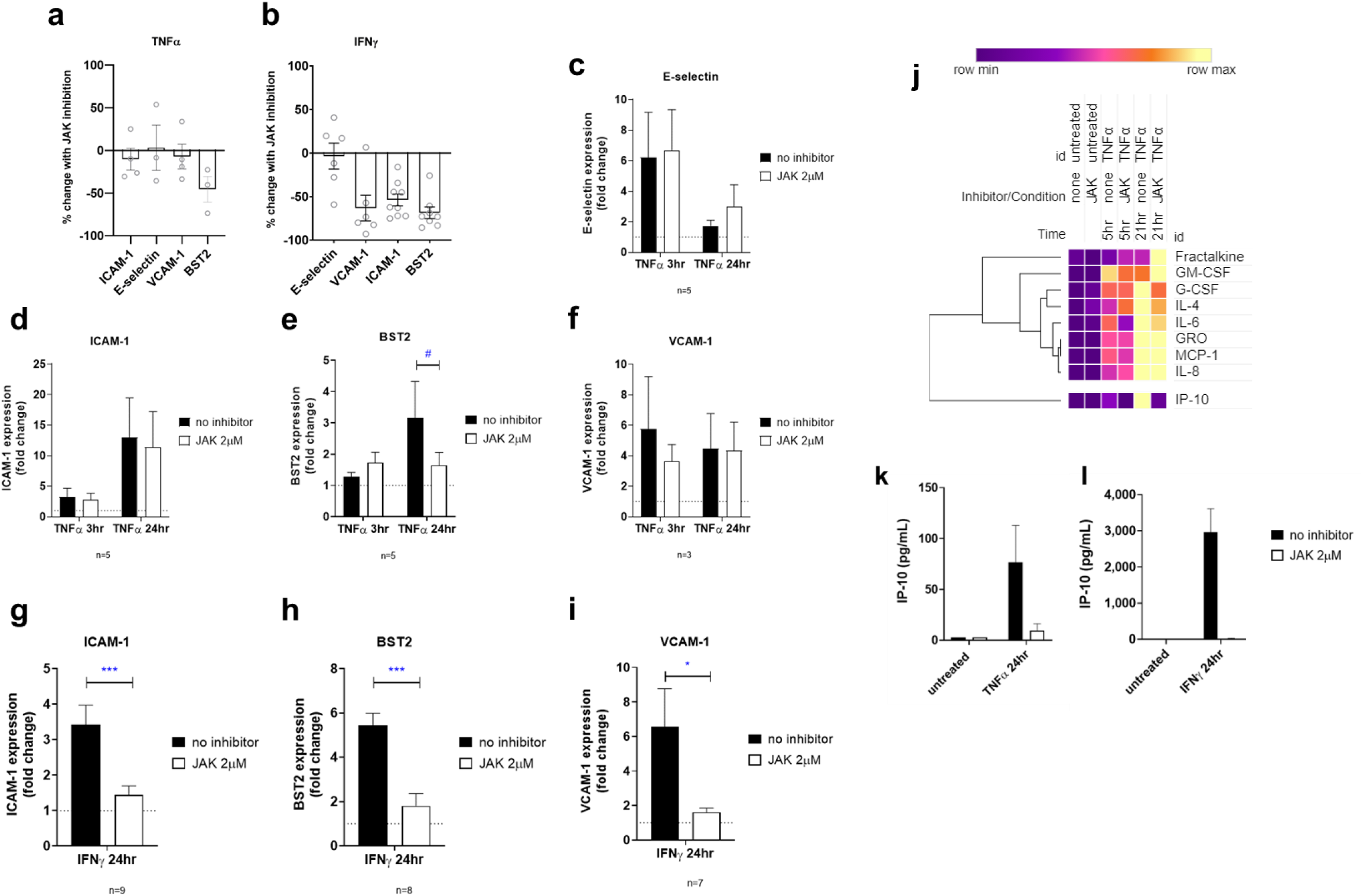
JAK-dependent endothelial cell activation. HAEC were pre-treated with the JAK1/2 inhibitor ruxolitinib (2μM) for 30min, prior to continuous stimulation with TNFα or IFNγ. Cell surface adhesion molecules were measured by flow cytometry, and secreted chemokines were measured by Luminex and ELISA. **a, b**. Graph shows percent inhibition by JAK inhibitor ruxolitinib of E-selectin (4hr), VCAM-1, ICAM-1 and BST2 (24hr) protein induction by HAEC in response to TNFα (panel a. n=3-5 independent experiments) or IFNγ (panel b, n=7-9 independent experiments). **c-f**. Fold change in TNFα-induced expression of E-selectin (c), ICAM-1 (d), BST2 (e), and VCAM-1 (f) without inhibitor (black bars) and with JAK inhibitor (white bars). Conditions were compared by t test, # p<0.1. n=3-5 independent experiments. **g-i**. Fold change in IFNγ-induced expression of ICAM-1 (g), BST2 (h) and VCAM-1 (i) without inhibitor (black bars) and with JAK inhibitor (white bars). Conditions were compared by t test, * p<0.05, *** p<0.001. n=7-9 independent experiments. **j**. HAEC were activated with TNFα or 5hr or 21hr. Heat map shows relative concentration of chemokines measured by Luminex, secreted by TNFα-stimulated HAEC, with and without JAK inhibition. **k, l**. IP-10 secretion by HAEC in response to continuous stimulation with TNFα (panel k, 24hr) or with IFNγ (panel l, 24hr), with and without JAK inhibition. Results are shown as mean concentration ±SEM (n=3 HAEC). Expression was compared to untreated endothelium by t test. * p<0.05; ** p<0.01; ***p<0.001, **** p<0.0001.

All IFNγ-stimulated adhesion molecules were significantly inhibited by pharmacological impairment of JAK: ICAM-1 (53.8±6.7%, **Figure 11b, 11g**), BST2 (68.5±6.7%, **Figure 11b, 11h**) and VCAM-1 (63.1±14.7%, **Figure 11b, 11i**). Raw concentration results for secreted IP-10 are shown in **Supplemental Figure 9g**.

The majority of TNFα-induced chemokines were unaffected by JAK inhibition (**Figure 11j**). At the mRNA level, JAK1/2 inhibition but not NFκB abolished *BST2* expression, as well as other STAT and interferon response genes (**Supplemental Figure 9h**). We analyzed the IFNγ and TNFα induced genes for overlap, stratifying those that were resolvable on TNFα withdrawal (defined as a difference of ≥2-fold higher mRNA counts in the continuous condition over the prime/rest condition) from those that were durable after short TNFα priming. 21 unique TNFα resolvable genes included NFκB genes and most chemokines like *NFKBIA, SELE, CXCL1, IL8, CCL20, CSF2, CCL2*, and *CCL5*; while 102 TNFα durable genes such as *STAT1, ICAM1*, and *BST2* were also IFNγ-inducible and JAK1/2 dependent (**Supplemental Figure 9i**).

Inhibition by ruxolitinib occurred in a dose-dependent manner (**Supplemental Figure 9j-9m**), with maximal inhibition at 2μM. Yet, IP-10 (*CXCL10*) expression in response to both TNFα (**Figure 11k**) and IFNγ (**Figure 11l**) was abrogated by JAK inhibition. Globally, we observed three patterns for TNFα-inducible genes: 1) sensitive only to NFκB inhibition (*CXCL8*/IL-8, *CCL2*/MCP-1, *CSF1*/M-CSF; *VCAM1, SELE*/E-selectin; *NFKBIA, NFKB1, RELB, TNFAIP3*); 2) partially sensitive to both NFκB and JAK inhibition (*CXCL10*/IP-10 (profound), IL-6 (partial), *ICAM1 (*partial*), CXCL1*/GROα (partial), *CXCL2*/MIP-2α (profound)); and 3) sensitive only to JAK inhibition (*BST2*). Among TNFα induced chemokines and adhesion molecules, only *BST2, CXCL10* (IP-10) and *CXCL2* (MIP-2α) were profoundly sensitive to JAK inhibition.

To examine possible crosstalk between *IRF1* and NFκB, we looked at TNFα-stimulated expression of NFκB or JAK inhibition. NFκB-related gene expressed was unaltered by JAK inhibition prior to TNFα treatment. When endothelial cells were pre-treated with MG132 to inhibit NFκB, TNFα induction of *IRF1* was unaffected. However, the JAK inhibitor blocked *IRF1* and *IRF9* transcription by TNFα. Interestingly, *STAT1* expression was blunted by either NFκB or JAK inhibition, suggesting parallel pathways.

### Durability of Intracellular Signaling after Cytokine Withdrawal

Given that ICAM-1, BST2 and IP-10 were sustained in the prime rest experiments, and that all three were JAK dependent, we compared whether NFκB or JAK/STAT gene expression were also differentially prolonged after cytokines were removed. There was a rapid decay of NFκB negative feedback regulators (*NFKBIA, NFKBIZ* and *TNFAIP3*) within 3hr of TNFα withdrawal (**Figure 12a-c**). This was followed by loss of enhanced expression of *NFKB1* (p105/p50), *NFKB2* (p100/p52), *RELA, RELB* and *BCL3* by 15hr after TNFα withdrawal (**Figure 12d-h**). Therefore in endothelial cells, NFκB gene expression appears to be closely aligned with TNFα presence.

**Figure 12.**
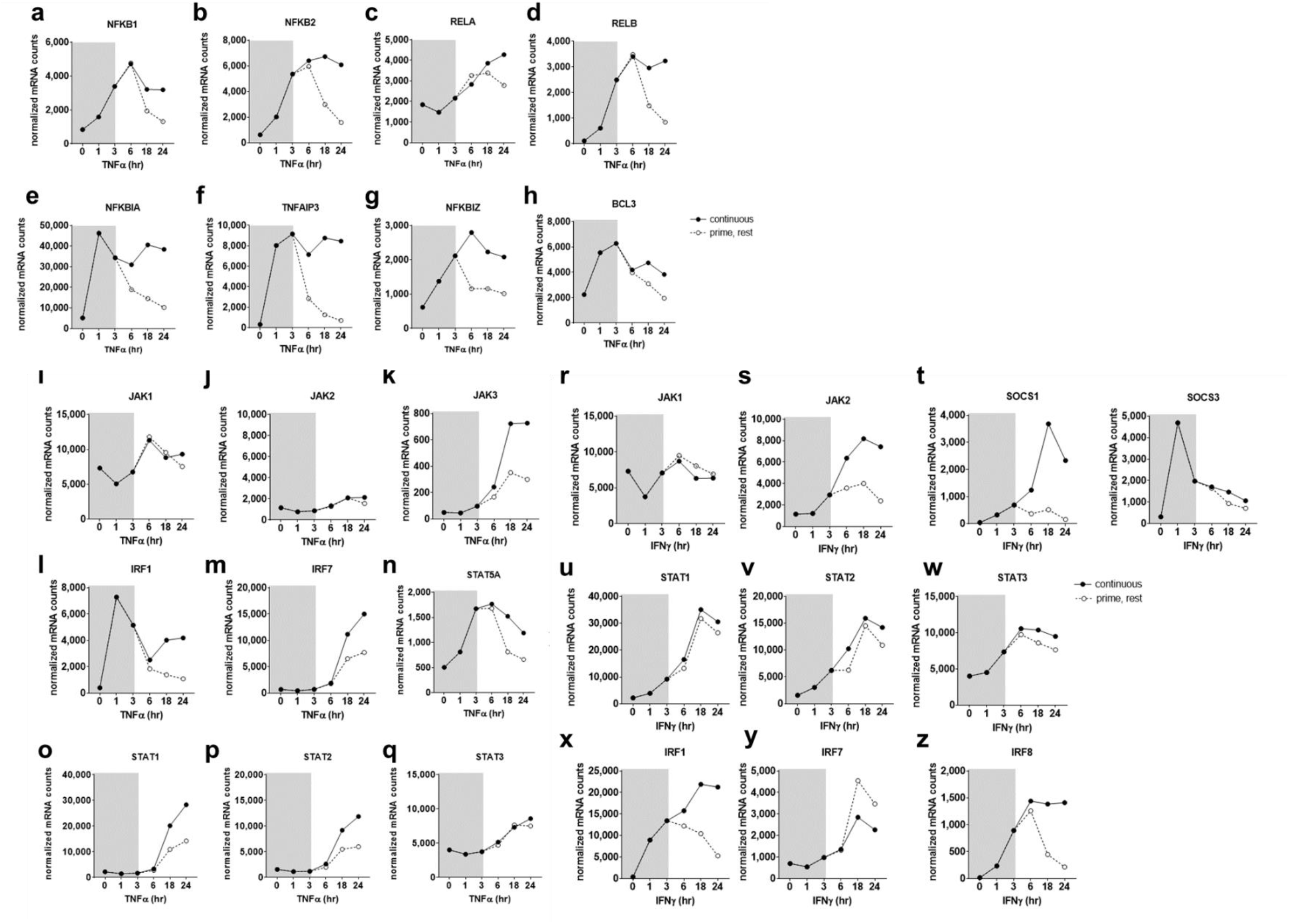
NFκB transcripts rapidly decline, and STAT transcripts persist, after withdrawal of TNFα and IFNγ. **a-h**. HMEC-1 were treated with TNFα for 1hr, 3hr, 6hr, 18hr or 24hr; or primed with TNFα for 3hr, and rested in normal medium. mRNA counts for NFκB related genes were measured by Nanostring. Results from one representative experiment are presented. The longitudinal levels of each transcript are shown under continuous TNFα stimulation (black circles, bold lines) or after prime/rest (open circles, dashed lines). **i-q**. HMEC-1 were treated with TNFα for 1hr, 3hr, 6hr, 18hr or 24hr; or primed with TNFα for 3hr, and rested in normal medium. mRNA counts for JAK, STAT, and interferon response genes were measured by Nanostring. Results from one representative experiment are presented. The longitudinal levels of each transcript are shown under continuous TNFα stimulation (black circles, bold lines) or after prime/rest (open circles, dashed lines). **r-z**. HMEC-1 were treated with IFNγ for 1hr, 3hr, 6hr, 18hr or 24hr; or primed with IFNγ for 3hr, and rested in normal medium. mRNA counts for JAK, STAT, and interferon response genes were measured by Nanostring. Results from one representative experiment are presented. The longitudinal levels of each transcript are shown under continuous TNFα stimulation (black circles, bold lines) or after prime/rest (open circles, dashed lines).

On the other hand, JAK/STAT gene expression was persistently elevated up to 24hr after withdrawal of either TNFα (**Figure 12i-p**). In particular, *JAK3, STAT1, STAT2* and *IRF7* mRNA continued to rise in TNFα-primed endothelial cells and remained elevated above baseline up to 21hr later.

Increased *JAK1, STAT1, STAT2* and *STAT3* expression was prolonged after IFNγ priming, while *JAK2* plateaued in the absence of continuous IFNγ (**Figure 12q-u**). Suppressor of cytokine signaling 1 (*SOCS1*) was not induced in the late phase after only short exposure to IFNγ (**Figure 12v**), while *SOCS3* was increased early and expression was lowly maintained (**Figure 12w**). Early increases in other downstream interferon response genes *IRF1, IFI16, IFI35* were also sustained through 24hr. While *IRF1* plateaued and *IRF8* declined when IFNγ was removed, *IRF7* transcript persisted even after IFNγ withdrawal. Hence, there was a maintenance of JAK/STAT transcripts after both TNFα and IFNγ stimulation, compared with rapid reduction in NFκB-related genes. We posited that this signaling was tied to the protracted expression of specific adhesion molecule/chemokine profiles prompted by IFNγ.

### Acute TNFα vs. chronic/desensitizing TNFα

We wanted to test our hypothesis that NFκB-driven inflammation is more readily resolvable than JAK/STAT mediated endothelial activation. Stimulation of endothelium with soluble TNFα (sTNFα) models an acute exposure, while a chronic exposure with stable transmembrane expression (tmTNFα) triggers desensitization to TNFα ^12,13^. We examined a dataset of murine endothelium stimulated with soluble/acute or chronic/desensitizing TNFα (GDS2773) with a particular focus on adhesion molecules, chemokines and NFκB vs. STAT-mediated transcription (**Figure 13a**). *Vcam1*, and chemokine transcripts *Csf1, Csf2*, and *Ccl2*, were not different between chronic and acute stimulation (**Figure 13b**). As reported ^13^, endothelial expression of *Sele* was greater with soluble/acute TNFα than chronic membrane bound TNFα. We also found that NFκB-dependent chemokines inlcuding *Cx3cl1, Cxcl1*, and *Cxcl12* were higher when EC were stimulated with soluble than tmTNFα (**Figure 13c**). Similarly, transcripts *Nfkb2, Rela*, and *Relb* were all significantly elevated with acute soluble TNFα but not chronic tmTNFα, as were the canonical NFκB target genes *Nfkbia, Nuak* and *Tfnaip3* (A20) (**Figure 13d**).

**Figure 13.**
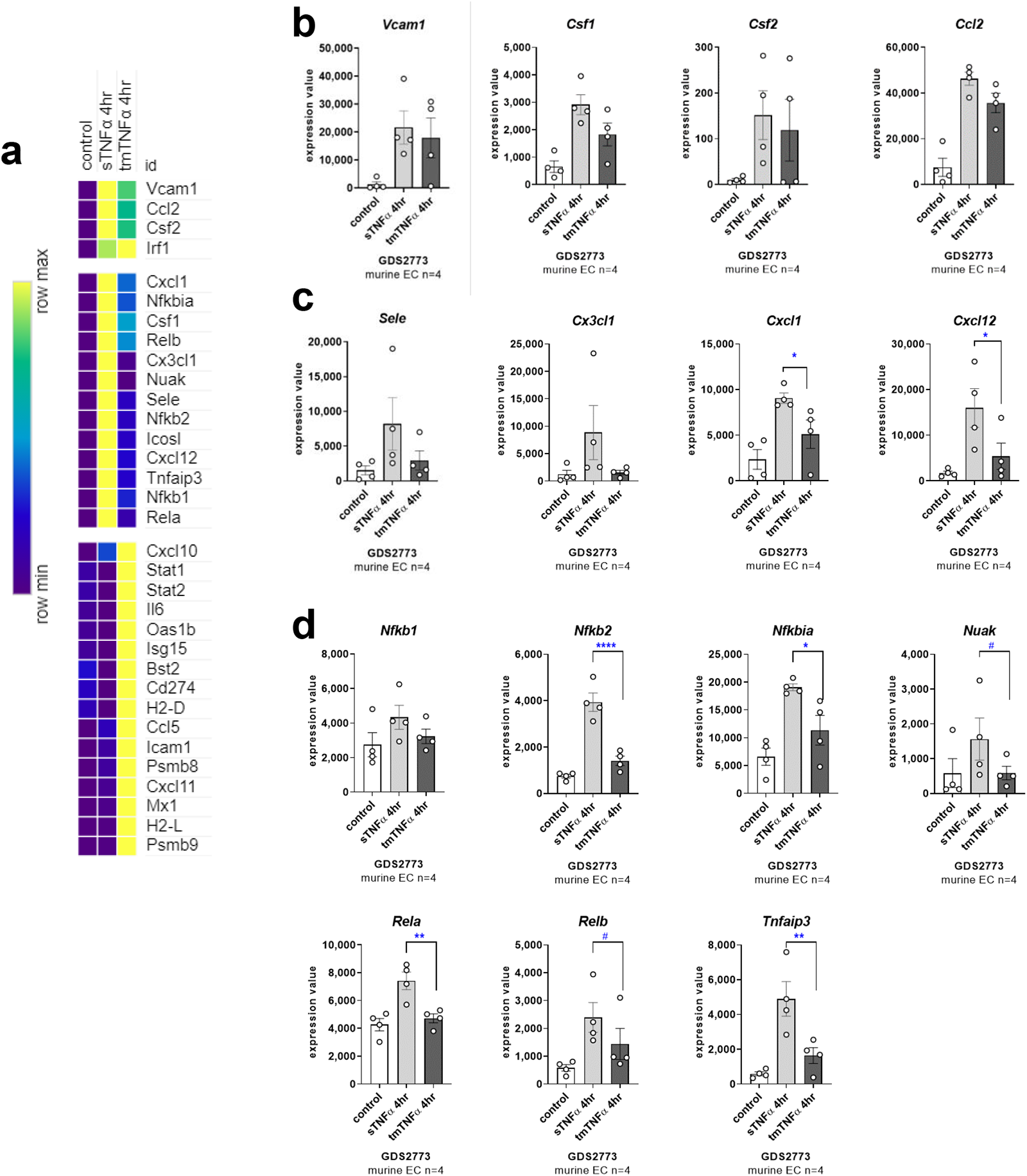
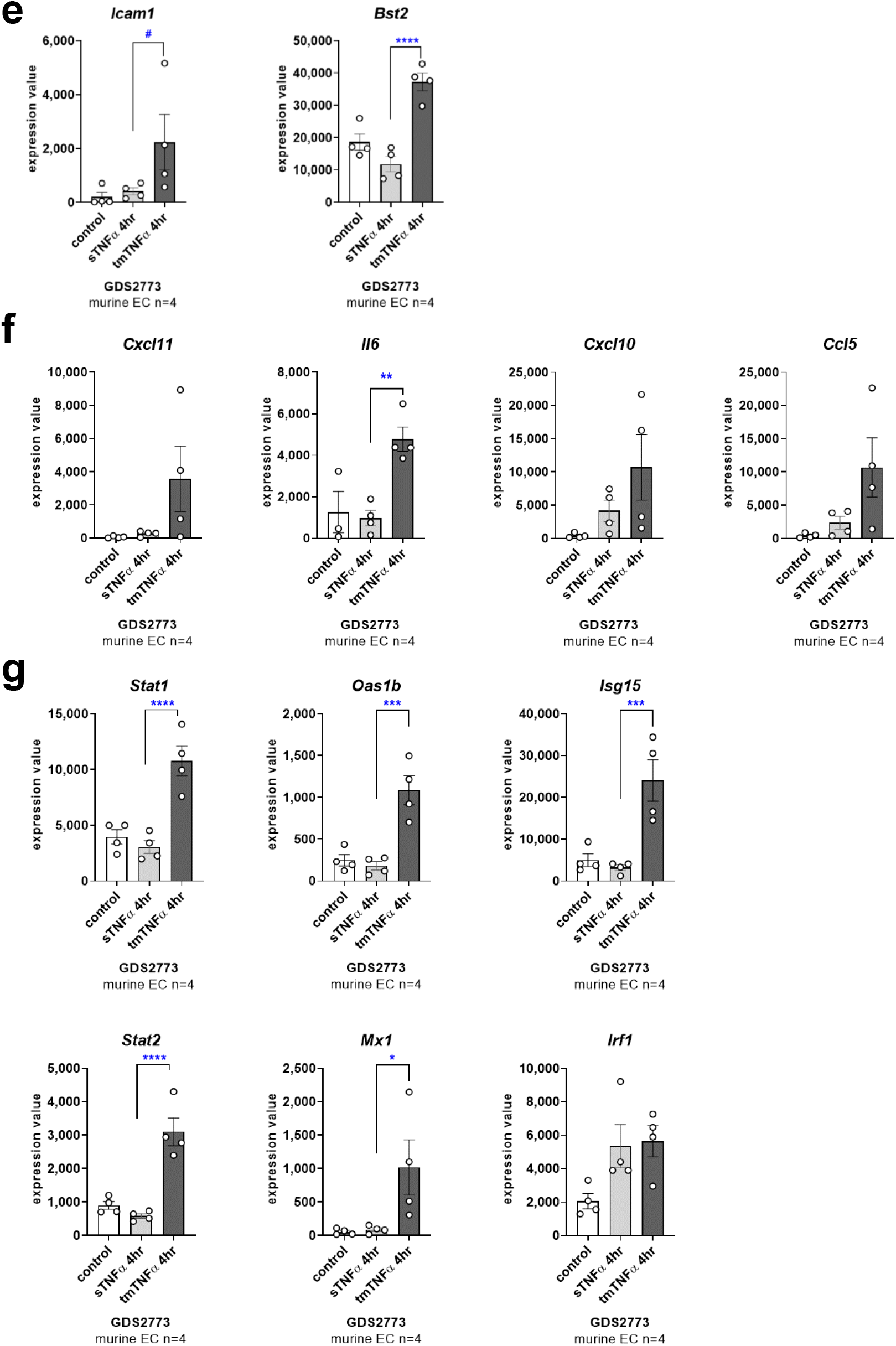
Endothelial cell gene expression is distinct after acute and chronic stimulation with TNFα. a) Heat map shows relative gene expression of adhesion molecules, chemokines, NFκB and interferon response genes in murine endothelial cells treated with acute, soluble TNFα (sTNFα) or chronic, desensitizing transmembrane TNFα (tmTNFα), from GDS2773. b) Expression values for genes *Vcam1, Csf1, Csf2, and Ccl2* which were not different comparing sTNFα to tmTNFα. c, d) Expression values for c) adhesion molecule *Sele* and chemokine *Cx3cl1, Cxcl1, Cxcl12* and d) NFκB related genes which were significantly higher with sTNFα than tmTNFα. Expression between groups by one way ANOVA followed by Fisher’s LSD. * p<0.05; ** p<0.01; ***p<0.001, **** p<0.0001. e, f) Expression values for e) adhesion molecules *Icam1, Bst2* and f) chemokines *Cxcl11, Il6, Cxcl10*, and *Ccl5* which were significantly higher with tmTNFα than sTNFα. g) Expression values for interferon response genes *Stat1, Stat2, Oas1b, Mx1, Isg15*, and *Irf1* which were significantly higher with tmTNFα than sTNFα

On the other hand, similar to our findings with TNFα withdrawal, *Icam1* and *Bst2* mRNA at 4hr were significantly higher with chronic, tmTNFα than soluble TNFα (**Figure 13e**). In addition, chemokines *Cxcl11, Cxcl10, Ccl5* and the cytokine *Il6* were significantly elevated with tmTNFα (**Figure 13f**). Unlike NFκB-dependent genes, STAT and interferon stimulated genes (ISGs) *Stat1, Stat2, Mx1, Oas1b*, and *Isg15* were only elevated with chronic TNFα, but not soluble TNFα (**Figure 13g**).

Taken together, these results support our findings that NFκB activation represents an acute, readily resolvable state, while JAK/STAT-driven endothelial inflammation (ICAM-1, BST2 and CXCL chemokines) is durable and chronic.

### JAK inhibition after inflammation initiation blunts the prolonged IFNγ response

Lastly, given that STAT transcription factor expression increased after TNFα and IFNγ priming, we examined whether inhibition of JAK/STAT after initial priming could prevent late upregulation of adhesion molecules. Endothelial cells were treated with TNFα or IFNγ for 3hr, then medium was replaced without IFNγ but with the JAK inhibitor ruxolitinib (2μM based on dosing studies **Supplemental Figure 8h-k**), the NFκB inhibitor MG132, or the transcription inhibitor CHX for an additional 21hr. ICAM-1 and BST2 on the cell surface were measured by flow cytometry, and IP-10 (*CXCL10*) and I-TAC (*CXCL11*) were measured in the supernatant by ELISA.

The presence of the NFκB inhibitor after TNFα priming did not block later upregulation of ICAM-1 (−10.1±1.3%, n=2 HAEC) or BST2 (42.1±39.9% change, n=2 HAEC). However, residual VCAM-1 expression was abrogated when NFκB was inhibited in the wash out phase (−58.7±5.19% decrease, n=2 HAEC). In contrast, late phase BST2 expression after TNFα priming was reduced in the presence of the JAK inhibitor ruxolitinib.

After initial priming with IFNγ for 3hr, late addition of a JAK1/2 inhibitor significantly prevented ICAM-1 (**Figure 14a**) and BST2 (**Figure 14b**) upregulation, and blocked IP-10 (**Figure 14c**) and I-TAC (**Figure 14d**) secretion, comparable to blocking all new transcription with CHX. Addition of the transcription inhibitor cycloheximide (CHX) in the rest period diminished the upregulation of ICAM-1, BST2, IP-10, and I-TAC, providing evidence that ongoing new transcription is required after IFNγ withdrawal. The same results were obtained using primary endothelial cells from four other vascular beds (**Figure 14e, 14f, 14g, 14h**). These results reveal that, while IFNγ presence is not required, active JAK signaling and ongoing new transcription are necessary for prolonged endothelial cell activation.

**Figure 14.**
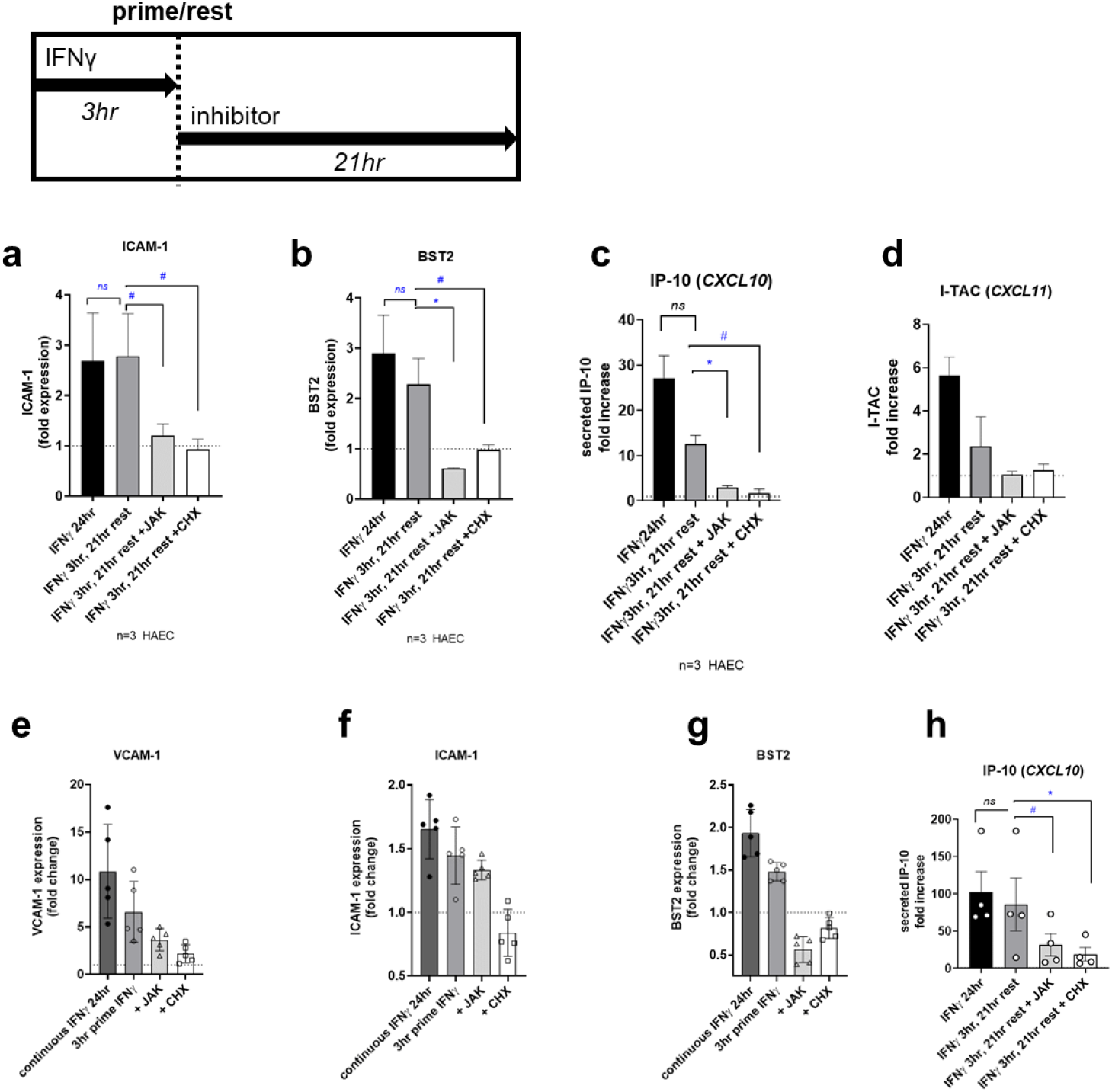
Pharmacological inhibition of JAK in the late phase prevents upregulation of ICAM-1, BST2, IP-10 and I-TAC. Endothelial cells were primed with IFNγ for 3hr, then medium was removed and replaced with normal medium or with medium containing the JAK1/2 inhibitor ruxolitinib (2μM) or the transcription inhibitor cycloheximide (CHX 10µM). Cell surface adhesion molecules were measured by flow cytometry. Secretion of chemokines into the supernatant was measured by ELISA. **a-b**. Expression of ICAM-1 (a) BST2 (b) on HAEC was measured by flow cytometry, after continuous IFNγ stimulation for 24hr (black bars), 3hr prime with IFNγ followed by 21hr rest (dark grey bars), 3hr prime with IFNγ followed by 21hr rest with JAK (light grey bars), or 3hr prime with IFNγ followed by 21hr rest with CHX (white bars). Groups were compared to uninhibited prime/rest condition by t test. # p<0.1, * p<0.05, ns not significant (n=3 HAEC). **c-d**. Secretion of IP-10 (c) and I-TAC (d) by HAEC was measured by ELISA, after continuous IFNγ stimulation for 24hr (black bars), 3hr prime with IFNγ followed by 21hr rest (dark grey bars), 3hr prime with IFNγ followed by 21hr rest with JAK (light grey bars), or 3hr prime with IFNγ followed by 21hr rest with CHX (white bars). Groups were compared to uninhibited prime/rest condition by t test. # p<0.1, * p<0.05, ns not significant (n=3 HAEC). **e-g**. Expression of VCAM-1 (e), ICAM-1 (f), and BST2 (g) on five primary endothelial cell types was measured by flow cytometry. Cells were exposed to continuous IFNγ stimulation for 24hr (black bars), 3hr prime with IFNγ followed by 21hr rest (dark grey bars), 3hr prime with IFNγ followed by 21hr rest with JAK (light grey bars), or 3hr prime with IFNγ followed by 21hr rest with CHX (white bars). **h**. Secretion of IP-10 by five primary endothelial cell types was measured by ELISA, after continuous IFNγ stimulation for 24hr (black bars), 3hr prime with IFNγ followed by 21hr rest (dark grey bars), 3hr prime with IFNγ followed by 21hr rest with JAK (light grey bars), or 3hr prime with IFNγ followed by 21hr rest with CHX (white bars). Results were compared to uninhibited prime/rest condition by t test. # p<0.1, * p<0.05, ns not significant.

## Discussion

In summary, our results demonstrate that short-term treatment with IFNγ elicits long-term signal propagation within the endothelial cell, in the absence of continuous cytokine stimulation, mediated by JAK1/2 and dependent on new transcription. On the other hand, TNFα elicits concurrent NFκB and JAK/STAT-dependent transcription, only the latter of which drives a durable, chronic inflammatory state.

Cytokines, reactive oxygen species, pathogen associated molecular patterns and endogenous danger signals cause type 2 activation of endothelium that is delayed but long lasting ^14^. Our results further illustrate the temporal phases of endothelial cell phenotype within the same continuous type II inflammatory stimulation, the likely biological consequence of which represents and acute/transient response vs. sustained chronic inflammatory phenotype. Our time course shows that endothelial cells exhibit sequential phases of activation by TNFα, with early and transient expression of the tethering adhesion molecule E-selectin and innate chemokines; intermediate, longer lasting expression of firm adhesion molecules ICAM-1 and VCAM-1 and adaptive chemokines; and finally late phase presentation of BST2 and CD164. Importantly, there are putative implications for the type of leukocyte recruited in these different phases that remain to be determined. Interestingly, the subset of mediators that persisted after TNFα withdrawal were ICAM-1, BST2, *CCL5* (RANTES), *CXCL10* (IP-10) and *CXCL11* (I-TAC), the latter of which are adaptive chemokines ^15^. In contrast, E-selectin, IL-8, *CCL2* (MCP-1), and *CCL20* (LARC/MIP-3α) rapidly resolved along with contraction of NFκB transcription, and predominantly recruit innate immune cells. It is tempting to speculate that the functional consequence is a shift from infiltration of the innate immune compartment including neutrophils and monocytes, to a more sustained recruitment of T cells.

Through successive phases of inflammation, recruited leukocytes may later secrete pro-resolution factors to counteract endothelial activation and switch to a resolving, wound healing environment ^3^; however, we observed that the acute effects of TNFα were transient, without the requirement for other cell types, suggesting cell intrinsic mechanisms shut down inflammation. Limited prior work has assessed the durability of endothelial activation after cytokine stimulation. TNFα and IL-1β induced changes in HLMVEC using a 1hr TNFα pulse followed by 3 and 19hr chase demonstrated that ICAM-1 did increase at 4hr, but VCAM-1, E-selectin, IL-6 and MCP-1 did not change with only 1hr priming ^16^. Similarly, in HUVEC treated with TNFα; ICAM-1 persisted at elevated levels at cell surface for up to 5 days ^17^. These early findings are in line with our results, which we elaborate on by describing the full repertoire of adhesion molecule and cytokine dynamics, as well as the differential transcriptional programming of intracellular signaling likely to be involved.

NFκB signaling is characterized by multi-phase responses, which can be associated with either reversible or long-term processes. Multiple potential mechanisms may govern the rapid contraction of NFκB-dependent functions (numerous endogenous and inducible negative feedback circuits). Recently, Sun et al. described discrete models of TNFα-induced mRNA dynamics, which were differentially regulated at the level of either chromatin-driven, transcriptional regulation or post-transcriptional mature mRNA half-life ^18^. Additional dynamics occur at the protein level; for example, TNFα induced E-selectin is subject internalization and rapid protein turn-over in addition to the short half-life of its mRNA ^19^. We posit that NFκB is reliant on direct, continuous receptor stimulation and signaling, while STAT activation is a secondary or self-sustaining response and therefore less quickly responsive to loss of stimulus. In part, NFκB may exhibit more negative feedback mechanisms compared with JAK/STAT, leading to rapid suppression once the balance shifts. Indeed, nuclear NFκB activity is transient following TNFα stimulation, limited in part by the actions of IκBα ^18^, which constitutes an early response negative feedback regulators. It was somewhat surprising that the endogenous feedback inhibitors of NFκB signaling, *NFKBIA, TNFAIP3*, and *IKBKE*, also so rapidly contracted, given that they would be expected to participate in resolution of NFκB signaling.

Collectively, these results demonstrate that IFNγ activation of endothelial cells results in predominantly late phase presentation of a restricted set of adhesion molecules and adaptive chemokines. One of the most robust adhesion molecules induced by IFNγ was BST2, which is much less well-characterized compared with ICAM-1, E-selectin and VCAM-1. Widely and constitutively expressed in the vasculature ^20^, BST2 is upregulated by interferons, possibly as an anti-viral host restriction factor ^21^, and mediated monocyte adhesion to endothelial cells ^22^.

These results also show differential regulation of endothelial adhesion molecules over time. ICAM-1 and BST2 induction required as little as 3hr of exposure to TNFα, and proceeded to increase to nearly the same extent after withdrawal as in the continuous presence of cytokine. However, the dynamics of E-selectin expression were consistent whether TNFα was continuously present or withdrawn. Moreover, VCAM-1 plateaued after short priming and declined when TNFα was removed. We provide preliminary evidence to suggest that NFκB dependent events rapidly contract, while JAK/STAT dependent events are more sustained. Of the TNFα-induced genes at 18hr that were JAK-dependent (eg. BST2, *CXCL10*), many were also among those that were persistently elevated after TNFα withdrawal; compared with the strongly NFκB-dependent genes (E-selectin, *CCL2*, VCAM-1, *TNFAIP3* etc) which required continuous TNFα for sustained expression.

Like NFκB, JAK/STAT pathways have several negative feedback regulators that blunt activation, perhaps best described are SOCS proteins SOCS1 and SOCS3. Recent evidence suggests that ISG expression is sustained, particularly of STAT1, STAT2, IRF1 and IRF9, due to a positive feedback loop ^11^. Our experiments offer additional proof of prolonged JAK/STAT dependent transcription after type II IFN, and reveal the functional consequences downstream.

Our results derive from *in vitro* experiments, and should be confirmed in the *in vivo* setting. Follow-up functional studies are also planned to link the persistent vs. resolvable endothelial pro-adhesive phenotypes with leukocyte recruitment. Here, we focused on EC adhesion molecules and chemokines involved in recruitment of immune cells. Endothelial cells are also reasonably efficient semi-professional antigen presenting cells ^23,24^. Although they lack B7 ligands needed to stimulate naïve T cells, endothelium expresses a host of other costimulatory and coinhibitory molecules and cytokines that shape the adaptive immune response ^23^. We recently showed that endothelial costimulatory phenotypes were distinct after TNFα versus IFNγ activation ^4^. We theorize that JAK/STAT signaling may also play a role in the enduring antigen presentation cell state of endothelial cells, a hypothesis that remains to be confirmed.

Organ transplant recipients must be maintained on life-long systemic immunosuppression to prevent rejection of the allograft, due to mismatched polymorphic antigens. Standard immunosuppression regimens rely heavily on calcineurin inhibitors and anti-metabolites to restrain activation and proliferation of recipient T cells. However, long-term use of systemic immunosuppression is associated with significant comorbidities and side effects, including increased rates of infection, new onset diabetes and malignancy post-transplant. Therefore there is a dire need for less intensive, more targeted therapies to suppress inflammation and alloimmune responses locally. Importantly, global transcript signatures of rejection in both the heart and the kidney are predominated by IFN-inducible genes, particularly the chemokines *CXCL9* (MIG), *CXCL10* (IP-10), *CXCL11* (I-TAC) and *CX3CL1* (fractalkine) ^25^. JAK3 inhibition has been tested in several clinical trials as an alternative immunosuppressant in solid organ transplantation [NCT00263328; NCT00483756; NCT00658359] and did show marked efficacy in preventing acute rejection. Although trials were withdrawn due to higher rates of viral infection and malignancy ^26,27^, there is continued interest in the potential application of other JAK inhibitors with improved safety profiles for transplant immunosuppression ^28,29^. JAK3 is preferentially expressed in T and NK cells, but not widely in the endothelium ^30^. New studies are evaluating JAK1/2 inhibitors in select SOTx recipient populations, such as those with carcinoma [NCT04807777] or with chronic bronchiolitis obliterans syndrome post-lung transplant [NCT03978637]. However, the effect of JAK antagonism on the donor allograft itself has yet to be considered.

Vessels from patients with giant cell arteritis had elevated expression of STAT1 and STAT2, and the JAK inhibitor tofacitinib reduced vasculitis and microangiogenesis in a mouse model ^31^. Tofacitinib also improved disease markers in lupus-prone mice, including endothelial function ^32^, as did a SOCS1 agonist peptide that represses JAK/STAT function ^33^. JAK/STAT responsive SOCS proteins were expressed within atherosclerotic lesions, and deficiency in SOCS worsened atherogenesis ^34^. Moreover, STAT1 signature in atherosclerotic plaques ^35^.

In conclusion, TNFα and IFNγ patterns of endothelial cell activation were distinct, where TNFα-inducible changes were mostly self-limiting over time, and the majority of which rapidly declined when TNFα was removed; compared with IFNγ whose effects were extended without the requirement for continual stimulation. Initiation of anti-TNFα therapy after inflammation has begun may resolve acute NFκB-mediated events but will likely not rapidly reverse the chronic phenotypic change. Our results indicate that JAKinibs may be beneficial in repressing chronic endothelial cell activation that leads to leukocyte infiltration into the tissue, in addition to its potential effects on the immune cell compartment.

## Materials and Methods

### Ethics Statement

Use of primary human endothelial cells for all experiments was approved by the UCLA Institutional Review Board (IRB#17-000477)

### Cells and Reagents

The list of sources for key biological materials and reagents can be found in **Supplemental Table 1**.

The immortalized human dermal microvascular cell line HMEC-1 was obtained from ATCC. Primary human aortic endothelial cells (HAEC) were obtained from commercial sources and aortic EC (HAEC) isolated from the aortic rings of deceased donors were kindly provided by Dr. EF Reed, UCLA. Primary human coronary artery (HCAEC), cardiac microvascular (HCMVEC), pulmonary artery (HPAEC), pulmonary microvascular (HPMVEC), dermal microvascular (HDMVEC), and renal glomerular (HRGEC) EC were obtained from commercial sources (**Supplemental Table 1**) and used between passages 3-8. All endothelial cells were cultured on 0.2% gelatin in complete endothelial cell growth medium (PromoCell). Experiments were carried out in M199+10% fetal bovine serum (FBS).

### Prime-Rest Experiments

Concentrations of TNFα and IFNγ were selected for maximal adhesion molecule protein induction based on initial titration experiments (**Supplemental Figure 1**). For transcript changes, endothelial cells (HMEC-1) and primary endothelial cells were stimulated with TNFα (20ng/mL) or IFNγ (200U/mL) diluted in M199+10% FBS for 1hr, 3hr, 6hr, 18hr and 24hr. For protein changes, primary endothelial cells (HAEC) were stimulated with TNFα (20ng/mL) or IFNγ (200U/mL) for 3hr-48hr. In prime-withdrawal conditions, endothelial cells were primed with TNFα or IFNγ for 3hr, then medium was removed, cells were washed, and fresh M199+10% FBS without cytokine was added for the remainder of the experiment. Negative control untreated cells were prepared at the same time points for comparison. Supernatants were collected for secreted analyte analysis along with matched mock treatment at each time point.

### Targeted Gene Expression Analysis

For transcript changes, cells were detached with trypsin, pelleted by centrifugation, and resuspended in RLT Buffer at 6,500 cells per μL (Qiagen). mRNA counts were measured by Nanostring (Human Immunology Panel). mRNA counts were measured by Nanostring (Human Immunology Panel v2.0, Nanostring Technologies) and analyzed in NCounter software. mRNA counts were normalized against internal and housekeeping controls. Normalized counts ≥250 were considered positive, and genes were considered changed if the counts differed by ±50% (i.e. 1.5-fold) or more compared with baseline.

### Public Dataset Analysis

Publicly available transcriptome data from GSE27870 (primary HUVEC, n=3, TNFα treatment 1hr, 1.5hr, 2hr, 3hr, 4hr, 5hr, 6hr); GDS2773 (murine endothelial cells stimulated with sTNFα or tmTNFα, 4hr, n=4); and GSE106524 (primary human lung microvascular endothelial cells, n=3-4, IFNγ treatment 3hr, 6hr, 24hr with paired mock time points) were analyzed GEO2R (https://www.ncbi.nlm.nih.gov/geo/info/geo2r.html) and graphed in Prism (GraphPad). Expression values were compared by two-way ANOVA followed by Fisher’s LSD, treated vs. mock for each group. We analyzed STAT1 Chip-Seq data from the ENCODE portal ^36,37^ with the following identifiers: ENCSR332EYT, ENCSR000EZK, ENCSR000EHK, ENCSR000EZK, ENCSR000EHJ, ENCSR000EGK, ENCSR000EGT, ENCSR000EZN. p65/RelA ChIP-Seq data in human endothelial cells ^38^ (GSE89970) was analyzed in the UCSC Genome Browser. The list of datasets can be found in **Supplemental Table 1**.

### Flow Cytometry, Cytokine and Chemokine Measurements

For protein changes, primary endothelial cells (HAEC) were stimulated with TNFα (20ng/mL) or IFNγ (200U/mL) diluted in M199 + 10% heat-inactivated FBS for 3hr, 6hr, 18hr, 24hr and 40hr. Cells were detached with Accutase, pelleted and resuspended in PBS + 2% heat-inactivated FBS. Cells were stained for ICAM-1-AF488, E-selectin-PE, VCAM-1-APC and BST2-PE/Cy7. Cell surface expression of adhesion molecule protein was measured by multiparameter flow cytometry (BD Fortessa). Supernatants were retained and secreted chemokines were measured by ELISA (IP-10, R&D Systems; I-TAC, R&D Systems) and/or Luminex (Milliplex Human 38-plex).

### Analyses

Technical replicates are defined as repeated measures of the same analyte with the same biological specimen. Biological replicates are defined as repeated measures of the same analyte with biological specimens from different sources and/or different experiments. With the exception of experiments employing the immortalized cell HMEC-1, experiments were repeated with a minimum of three biological replicates, as indicated in the figure legends. Outliers were not excluded from analyses except where negative or positive control conditions failed. Heat maps and hierarchical clustering were generated using Morpheus (https://software.broadinstitute.org/morpheus/). Source data for heat maps can be found in **Supplemental Table 2**. Venn diagram was generated in InteractiVenn ^39^. Graphs and statistical analyses were performed in Prism (GraphPad, San Diego, CA).

## Acknowledgements

The author would like to acknowledge the ENCODE Consortium and ENCODE production laboratory for generating the ChIP-Seq dataset analyzed herein; and thank Dr. Maura Rossetti (formerly UCLA) and the UCLA Immune Assessment Core for guidance on multicolor flow cytometry panels, Gemalene Sunga (UCLA) for technical support in conducting Luminex assays, Hasitha Gunawardana (UCLA) and Gianna Zufall for technical support in acquisition of some ELISA and flow data, and Dr. Emmanuelle Faure at the UCLA IMT Core/Center for Systems Biomedicine, which is supported by CURE/P30 DK041301, for conducting Nanostring services and providing technical assistance.

## Author Contributions

Experiment conception and design; data collection and formal analysis; manuscript preparation: NMV.

## List of Supplemental Materials

Supplemental Figures 1-9

Supplemental Table 1. Key Resources

Supplemental Table 2. Source Data for Heat Maps

