## Supplemental Table 1-1 for "Endothelial Resolution of Inflammation is Delayed Following JAK-driven but not NFκB-dependent Activation"

| Reagent type (species) or resource | Designation | Source or refe | Identifiers | Additional Information |
|---|---|---|---|---|
| Recombinant protein | TNF alpha (recombinant, Humankine) | Sigma | H8916-10UG | |
| Recombinant protein | Recombinant human IFNγ protein | R&D Systems | 285IF-100/CF | |
| Antibody | ICAM-1-Alexa Fluor 488 | Biolegend | 322714 | |
| Antibody | E-selectin-PE | Biolegend | 322606 | |
| Antibody | VCAM-1-APC | Biolegend | 305810 | |
| Antibody | BST2-PE/Cy7 | Biolegend | 348416 | |
| Chemical compound, drug | In solution JAK I Inhibitor | Millipore Sigma | 420097-500UG | |
| Chemical compound, drug | Ruxolitinib | Invivogen | tlrl-rux | |
| Chemical compound, drug | MG132 | Selleck Chemi | S2619 | |
| Commercial assay or kit | CXCL11/I-TAC DuoSet ELISA | R&D | DY672 | |
| Commercial assay or kit | CXCL10/IP-10 DuoSet ELISA | R&D | DY266-05 | |
| Commercial assay or kit | CCL20/MIP3a DuoSet ELISA | R&D | DY360-05 | |
| Commercial assay or kit | DuoSet ELISA Ancillary Reagent Kit 2 | R&D | DY008 | |
| Commercial assay or kit | Milliplex MAP Human Cytokine/Chemokine 38-Plex panel | Millipore Sigma | HCYTMAG-60K-PX38 | |
| Cell line (*Homo sapiens*) | HMEC-1 | ATCC | CRL-3243 | Immortalized cell line |
| Cell line (*Homo sapiens*) | HAEC (human aortic endothelial cells) | PromoCell | C-12271 | Primary cells |
| Cell line (*Homo sapiens*) | HAEC (human aortic endothelial cells) | Lonza | CC-2535 | Primary cells |
| Cell line (*Homo sapiens*) | HPAEC (human pulmonary artery endothelial cells) | PromoCell | CC-12241 | Primary cells |
| Cell line (*Homo sapiens*) | HCAEC (human coronary artery endothelial cells) | PromoCell | CC-12221 | Primary cells |
| Cell line (*Homo sapiens*) | HCMVEC (human cardiac microvascular endothelial cells) | PromoCell | C-12285 | Primary cells |
| Cell line (*Homo sapiens*) | HLMVEC (human lung microvascular endothelial cells) | PromoCell | C-12281 | Primary cells |
| Cell line (*Homo sapiens*) | HRGEC (human renal glomerular endothelial cells) | ScienCell | 4000 | Primary cells |
| Cell line (*Homo sapiens*) | HDBEC-c (blood dermal, abdominal skin endothelial cells) | PromoCell | C-12225 | Primary cells |
| Other | Trypsin EDTA | Sigma | T4049-100mL | |
| Other | Accutase | Sigma | A6964-500ML | |
| Other | Endothelial Cell Growth Medium MV | PromoCell | C-22020 | |
| Other | M199, Gibco | ThermoFisher | 11150059 | |
| Other | Fetal bovine serum | R&D Systems | S11150 | Heat-inactivated prior to use |
| Other | gelatin type B solution | Sigma | G1393-100ML | |
| Other | Buffer RLT | Qiagen | 79216 | |

**Supplemental Table 1. Key Resources and Reagents**
