## Supplemental Table 1-2 for "Endothelial Resolution of Inflammation is Delayed Following JAK-driven but not NFκB-dependent Activation"

| Data Avail | Accession | Description | Cell Type | n | Stimulus | Time Points | Reference |
|---|---|---|---|---|---|---|---|
| GEO | GSE27870 | Time course of TNFα stimulation of endothelium | HUVEC | n=3 | TNFα | 1-6hr | |
| GEO | GSE106524 | Time course of IFNγ stimulation of endothelium | HLMVEC | n-4 | IFNγ | 3-24hr | |
| GEO | GSE31477 | Transcription Factor Binding Sites by ChIP-seq | | | | | ENCODE |
| GEO | GSE89970 | p65/RelA ChIP-Seq on HAEC stimulated with TNFa for 4hr | HAEC | n=2 | TNFα | 4hr | PMID 28585919 |
| ENCODE | ENCSR332EYT | STAT1 ChIP-seq on human GM12878 | GM12878 | | | | |
| ENCODE | ENCSR000EZK | STAT1 ChIP-seq on human HeLa-S3 treated with IFNg for 30 minutes | HeLa | | | | |
| ENCODE | ENCSR000EHK | STAT1 ChIP-seq on human K562 treated with IFNg30 | K562 | | | | |
| ENCODE | ENCSR000EZK | STAT1 ChIP-seq on human HeLa-S3 treated with IFNg for 30 minutes | HeLa | | | | |
| ENCODE | ENCSR000EHJ | STAT1 ChIP-seq on human K562 treated with IFNg6h | K562 | | | | |
| ENCODE | ENCSR000EGK | IRF1 ChIP-seq on human K562 treated with IFNg30 produced by the Snyder lab | K562 | | | | |
| ENCODE | ENCSR000EGT | IRF1 ChIP-seq on human K562 treated with IFNg6h | K562 | | | | |
| ENCODE | ENCSR000EZN | Control ChIP-seq on human HeLa-S3 treated with IFNg for 30 minutes | HeLa | | | | |

**Supplemental Table 1. Key Resources and Reagents**
