## Supplemental Table 2 for "Endothelial Resolution of Inflammation is Delayed Following JAK-driven but not NFκB-dependent Activation"

**Supplemental Table 2. Source Data for Heat Maps--Related to Figure 1**

*normalized mRNA counts*

| | UT | TNFα 1hr | TNFα 3hr | TNFα 6hr | TNFα 18hr | TNFα 24hr | IFNγ 1hr | IFNγ 3hr | IFNγ 6hr | IFNγ 18hr | IFNγ 24hr |
|---|---|---|---|---|---|---|---|---|---|---|---|
| CCL2 | 198.21 | 9064.56 | 12094.07 | 16638.97 | 34106.28 | 30390.52 | 1484.35 | 733.08 | 451.15 | 562.21 | 462.27 |
| CCL20 | 6.24 | 1286.48 | 2733.65 | 2339.22 | 2976.47 | 2593.03 | 18.28 | 5.43 | 8.30 | 31.10 | 29.70 |
| CCL5 | 61.68 | 97.92 | 1587.57 | 1739.97 | 8196.58 | 8641.68 | 32.90 | 52.22 | 104.25 | 253.59 | 267.29 |
| CSF1 | 171.18 | 752.89 | 1204.06 | 1507.24 | 2025.11 | 2147.15 | 599.59 | 1308.82 | 1628.38 | 1445.01 | 1399.73 |
| CSF2 | 8.32 | 3029.73 | 1208.51 | 1210.38 | 597.33 | 229.58 | 14.62 | 4.75 | 7.38 | 11.96 | 5.17 |
| CXCL1 | 1489.34 | 37229.84 | 26939.62 | 21075.19 | 26794.05 | 15891.29 | 1020.04 | 521.50 | 649.51 | 1495.25 | 1297.72 |
| CXCL10 | 10.40 | 21.70 | 165.00 | 469.43 | 10749.09 | 13692.39 | 102.37 | 1402.41 | 1942.06 | 5748.92 | 4901.64 |
| CXCL11 | 11.78 | 13.49 | 80.27 | 187.61 | 5489.64 | 8305.94 | 233.99 | 2984.53 | 6936.06 | 26388.12 | 28131.50 |
| CXCL2 | 302.16 | 9713.66 | 7206.49 | 8042.02 | 9856.00 | 4050.12 | 431.41 | 360.10 | 563.71 | 1232.08 | 936.17 |
| CXCL9 | 6.93 | 5.86 | 13.38 | 8.71 | 68.47 | 90.24 | 29.25 | 539.81 | 1313.77 | 4830.25 | 4940.38 |
| IL8 | 1233.61 | 29668.13 | 71864.27 | 77746.95 | 89929.28 | 70962.07 | 1093.16 | 593.38 | 580.31 | 708.15 | 862.56 |
| ICAM1 | 624.43 | 28812.63 | 63172.77 | 52094.65 | 36338.27 | 32471.31 | 3612.17 | 5358.04 | 6636.22 | 7122.16 | 6142.55 |
| SELE | 5.54 | 189.98 | 3366.90 | 1007.73 | 67.02 | 72.99 | 3.66 | 6.10 | 8.30 | 19.14 | 7.75 |
| VCAM1 | 16.63 | 412.80 | 8321.36 | 13946.69 | 9612.70 | 7424.78 | 3.66 | 52.90 | 91.34 | 210.53 | 331.86 |
| BST2 | 32.57 | 14.66 | 8.92 | 63.33 | 3085.74 | 6355.19 | 7.31 | 122.74 | 381.95 | 4335.02 | 4026.16 |
| CD164 | 5824.98 | 4568.93 | 4223.11 | 6108.11 | 10995.31 | 13963.10 | 4745.54 | 6051.11 | 6617.77 | 7083.88 | 7105.83 |
| CD44 | 6277.53 | 4765.95 | 8829.74 | 11962.11 | 15894.90 | 15794.42 | 3842.50 | 4527.99 | 5903.68 | 8914.06 | 8226.65 |
| CD97 | 999.36 | 577.57 | 419.19 | 577.09 | 1277.71 | 1431.87 | 467.97 | 661.87 | 939.20 | 2045.50 | 2069.90 |

**Supplemental Table 2. Source Data for Heat Maps--Related to Figure 9a**

*normalized mRNA counts*

| | UT | IFNγ 1hr | IFNγ 3hr | IFNγ 6hr | IFNγ 18hr | IFNγ 24hr | TNFα 1hr | TNFα 3hr | TNFα 6hr | TNFα 18hr | TNFα 24hr |
|---|---|---|---|---|---|---|---|---|---|---|---|
| BCL3 | 1190.64 | 2910.21 | 2772.95 | 1572.10 | 2966.57 | 2299.74 | 5551.08 | 6278.93 | 4190.02 | 4755.36 | 3832.49 |
| IKBKAP | 1509.44 | 701.96 | 846.33 | 1051.76 | 1523.96 | 1380.36 | 727.09 | 784.87 | 824.86 | 1719.15 | 1628.28 |
| IKBKE | 675.71 | 321.73 | 406.21 | 614.45 | 1423.47 | 1176.34 | 442.12 | 1007.84 | 2223.65 | 4653.37 | 4757.43 |
| IKBKG | 1535.77 | 804.33 | 1093.85 | 1376.51 | 1933.05 | 1944.64 | 866.64 | 874.06 | 1251.54 | 2019.28 | 2145.82 |
| NFKB1 | 733.93 | 515.50 | 775.80 | 842.33 | 1016.77 | 951.66 | 1583.76 | 3384.73 | 4708.53 | 3208.12 | 3184.89 |
| NFKB2 | 458.79 | 416.79 | 458.43 | 509.27 | 760.78 | 746.35 | 2012.98 | 5346.90 | 6398.63 | 6722.19 | 6079.16 |
| NFKBIA | 5047.39 | 5769.24 | 4043.79 | 4310.36 | 5717.82 | 6847.58 | 46251.02 | 34328.96 | 30973.55 | 40590.99 | 38383.27 |
| NFKBIZ | 410.97 | 204.74 | 646.95 | 776.82 | 858.87 | 959.41 | 1375.02 | 2113.79 | 2794.40 | 2227.62 | 2080.80 |
| RELA | 1846.25 | 1239.40 | 1715.71 | 2084.14 | 2839.77 | 2711.66 | 1475.29 | 2158.38 | 2829.23 | 3854.99 | 4273.06 |
| RELB | 115.04 | 149.90 | 228.54 | 211.27 | 229.67 | 242.76 | 610.40 | 2492.84 | 3403.95 | 2958.99 | 3232.67 |
| TNFAIP3 | 252.27 | 91.40 | 173.61 | 202.97 | 354.07 | 494.55 | 8028.46 | 9155.28 | 7140.37 | 8753.12 | 8457.22 |

**Supplemental Table 2. Source Data for Heat Maps--Related to Figure 10a**

*normalized mRNA counts*

| | UT | IFNγ 1hr | IFNγ 3hr | IFNγ 6hr | IFNγ 18hr | IFNγ 24hr | TNFα 1hr | TNFα 3hr | TNFα 6hr | TNFα 18hr | TNFα 24hr |
|---|---|---|---|---|---|---|---|---|---|---|---|
| IRF1 | 366.37 | 8939.03 | 13425.97 | 15740.38 | 21904.77 | 21258.09 | 7279.68 | 5155.14 | 2514.96 | 4015.25 | 4180.17 |
| IRF3 | 493.00 | 288.83 | 406.21 | 473.29 | 1172.27 | 947.79 | 354.75 | 214.05 | 375.23 | 715.34 | 654.23 |
| IRF7 | 695.03 | 537.44 | 975.18 | 1357.14 | 2846.95 | 2264.88 | 452.67 | 700.14 | 1864.26 | 11149.74 | 14990.23 |
| IRF8 | 14.94 | 233.99 | 893.80 | 1440.17 | 1385.20 | 1410.06 | 14.07 | 35.68 | 10.29 | 50.99 | 91.57 |
| JAK1 | 7314.56 | 3743.79 | 7064.94 | 8711.13 | 6315.92 | 6355.61 | 5066.16 | 6760.55 | 11308.23 | 8818.68 | 9311.83 |
| JAK2 | 1154.59 | 1217.46 | 2948.59 | 6362.21 | 8179.60 | 7428.65 | 776.93 | 869.60 | 1346.54 | 2092.12 | 2139.19 |
| JAK3 | 49.80 | 40.22 | 75.95 | 61.81 | 107.66 | 87.81 | 46.32 | 98.11 | 243.03 | 722.63 | 727.22 |
| STAT1 | 2190.38 | 3882.72 | 9177.37 | 16498.75 | 35070.11 | 30524.21 | 1423.69 | 1658.92 | 3298.66 | 20115.57 | 28269.91 |
| STAT2 | 1564.36 | 3030.86 | 6196.23 | 10233.41 | 15902.25 | 14206.50 | 1122.88 | 1177.30 | 2611.54 | 9175.62 | 11850.46 |
| STAT3 | 4012.98 | 4507.90 | 7357.22 | 10568.32 | 10378.21 | 9493.38 | 3384.48 | 3750.41 | 5158.17 | 7312.24 | 8579.31 |
| STAT5A | 502.96 | 307.11 | 453.68 | 479.75 | 954.57 | 941.33 | 810.35 | 1672.30 | 1766.89 | 1521.01 | 1187.70 |

**Supplemental Table 2. Source Data for Heat Maps--Related to Figure 11j**

*pg/mL secreted protein*

| Stimulation | Time | Inhibitor/Cc | G-CSF | GM-CSF | Fractalkine | GRO | IL-4 | IL-6 | IL-8 | IP-10 | MCP-1 |
|---|---|---|---|---|---|---|---|---|---|---|---|
| untreated | 5hr | none | 7.25 | 1.45 | 19.6 | 110.62 | 0.06 | 17.13 | 282.78 | 2.89 | 341.4 |
| untreated | 5hr | JAK | 9.26 | 1.45 | 14.03 | 114.28 | 3.69 | 22.5 | 286.41 | 2.89 | 296.78 |
| TNFα | 5hr | none | 18.61 | 22.28 | 25.08 | 606.45 | 7.49 | 76.84 | 2139 | 40.35 | 5478 |
| TNFα | 5hr | JAK | 18.61 | 17.94 | 35.69 | 572.21 | 11.36 | 50.01 | 2429 | 3.2 | 4526 |
| TNFα | 21hr | none | 24.63 | 18.43 | 35.69 | 1004 | 16.26 | 108.66 | 4230 | 112.83 | 9030 |
| TNFα | 21hr | JAK | 19.28 | 25.06 | 64.53 | 952.13 | 13.31 | 97.27 | 4263 | 16.23 | 8596 |

**Supplemental Table 2. Source Data for Heat Maps--Related to Figure 13a**

*normalized mRNA expression values, mean of n=4*

| | control | sTNFα 4hr | tmTNFα 4hr |
|---|---|---|---|
| Vcam1 | 1306.20 | 21620.75 | 17876.35 |
| Sele | 1514.25 | 8199.25 | 2923.75 |
| Icam1 | 202.10 | 411.28 | 2232.50 |
| Cxcl1 | 2350.50 | 9061.50 | 5116.00 |
| Ccl2 | 7491.25 | 46294.00 | 35660.50 |
| Cxcl10 | 390.00 | 4139.25 | 10673.75 |
| Bst2 | 18628.50 | 11821.00 | 37267.75 |
| Ccl5 | 415.00 | 2333.25 | 10655.50 |
| Cxcl11 | 65.25 | 269.50 | 3568.25 |
| Cx3cl1 | 1230.75 | 8854.75 | 1564.75 |
| Il6 | 1256.67 | 968.25 | 4769.00 |
| Csf2 | 9.65 | 151.45 | 118.75 |
| Csf1 | 656.18 | 2914.75 | 1825.75 |
| Stat1 | 3962.25 | 3053.00 | 10769.75 |
| Stat2 | 898.50 | 576.75 | 3103.50 |
| Irf1 | 2058.00 | 5353.75 | 5646.00 |
| Cxcl12 | 1731.50 | 15981.00 | 5385.25 |
| Mx1 | 54.90 | 86.83 | 1016.15 |
| Oas1b | 245.53 | 180.85 | 1084.00 |
| Isg15 | 4967.25 | 3180.00 | 24028.50 |
| Nfkb1 | 2750.00 | 4334.25 | 3232.00 |
| Nfkb2 | 748.25 | 3935.00 | 1409.75 |
| Rela | 4267.25 | 7416.50 | 4714.25 |
| Relb | 583.00 | 2396.75 | 1440.75 |
| Nfkbia | 6594.25 | 19070.75 | 11350.00 |
| Tnfaip3 | 590.50 | 4895.25 | 1641.50 |
| Nuak | 584.25 | 1563.75 | 588.25 |
| Cd274 | 311.25 | 200.25 | 727.50 |
| Icosl | 80.50 | 549.43 | 185.25 |
| H2-L | 4336.50 | 4449.50 | 16791.25 |
| H2-D | 8243.75 | 5973.75 | 18785.00 |
| Psmb8 | 3077.75 | 3844.50 | 10904.50 |
| Psmb9 | 1074.00 | 1108.25 | 4695.75 |
