## Supplemental Figures for "Endothelial Resolution of Inflammation is Delayed Following JAK-driven but not NFκB-dependent Activation"

Nicole M Valenzuela

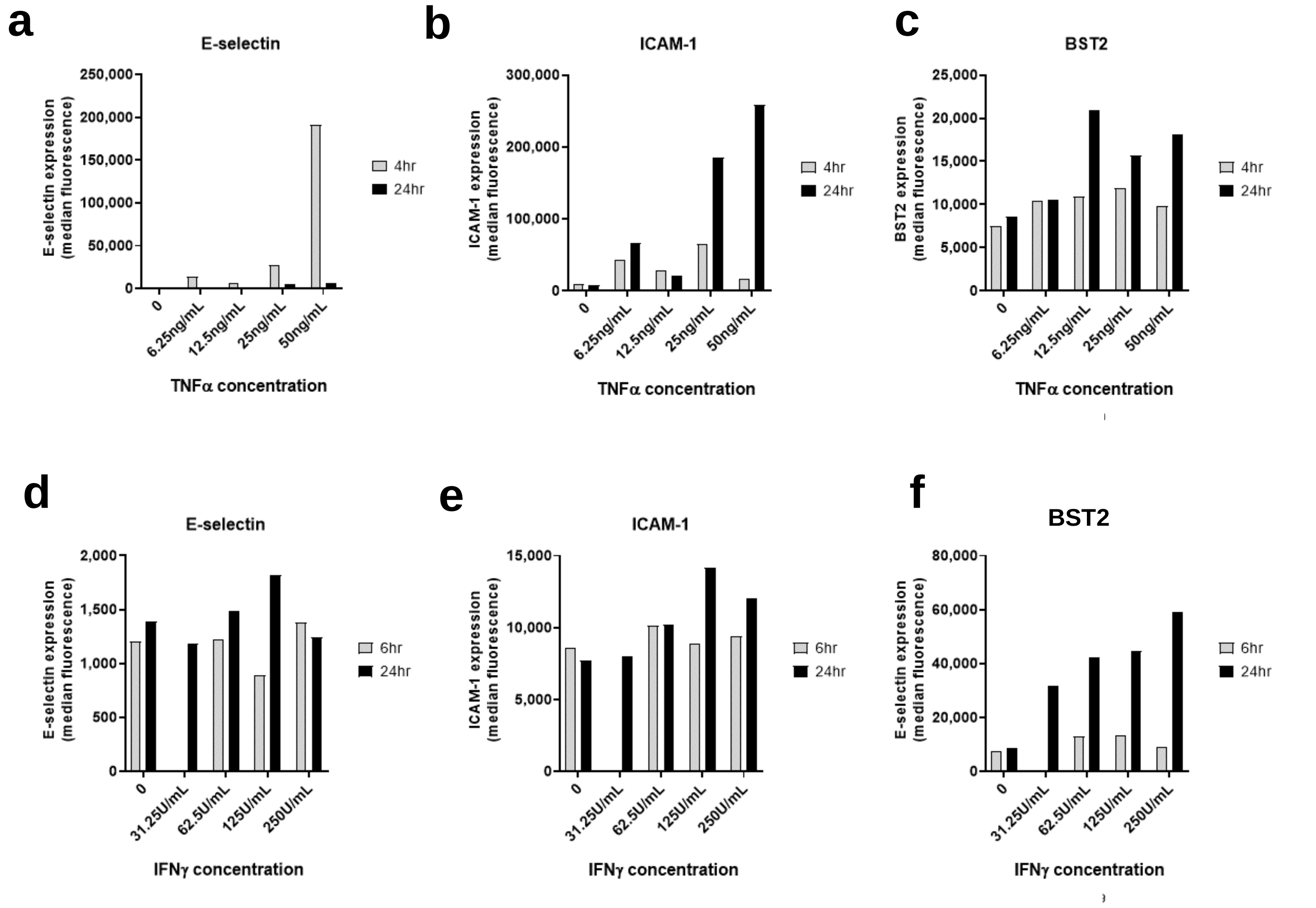


**Supplemental Figure 1. Preliminary dose titration studies.**
**a-c.** Primary human aortic endothelial cells (HAEC) were stimulated with TNFα (6.25-50ng/mL) for 4hr or 24hr. Cell surface E-selectin (a), VCAM-1 (b), ICAM-1 (c) were measured by multiparameter flow cytometry.
**d-f.** Primary human aortic endothelial cells (HAEC) were stimulated with IFNγ (31.25-250U/mL) for 6hr or 24hr. Cell surface E-selectin (d), VCAM-1 (e), ICAM-1 (f) were measured by multiparameter flow cytometry.

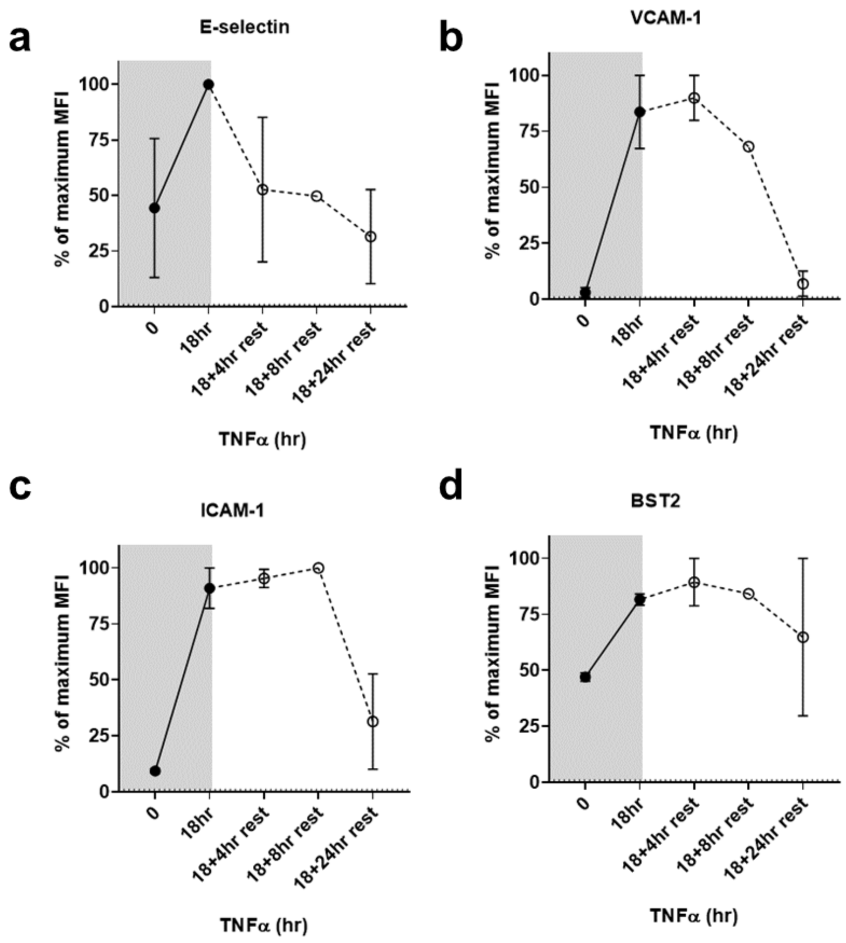


**Supplemental Figure 2.** Persistence of cell surface adhesion molecules after priming with TNFα, followed by cytokine withdrawal.

**a-d**. Human aortic endothelial cells were treated with TNFα for 18hr, then medium was removed and cells were rested for an additional 4hr, 8hr or 24hr in normal medium (n=2). Cell surface E-selectin (a), VCAM-1 (b), ICAM-1 (c), and BST2 (d) were measured by multiparameter flow cytometry. Results are expressed as the percent of maximum MFI within each experiment.

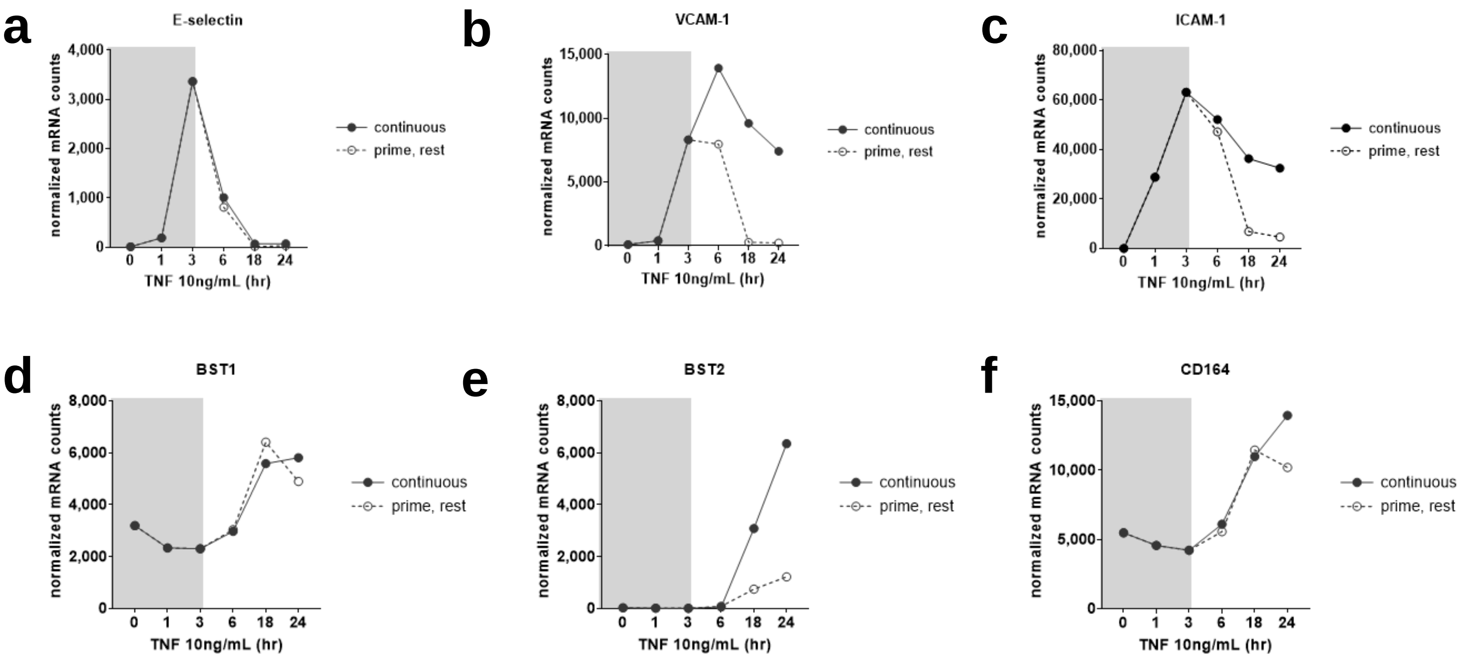


**Supplemental Figure 3.** HMEC-1 were treated with TNFα, followed by cytokine withdrawal for 3hr, 15hr or 21hr. Normalized mRNA counts were measured by Nanostring.

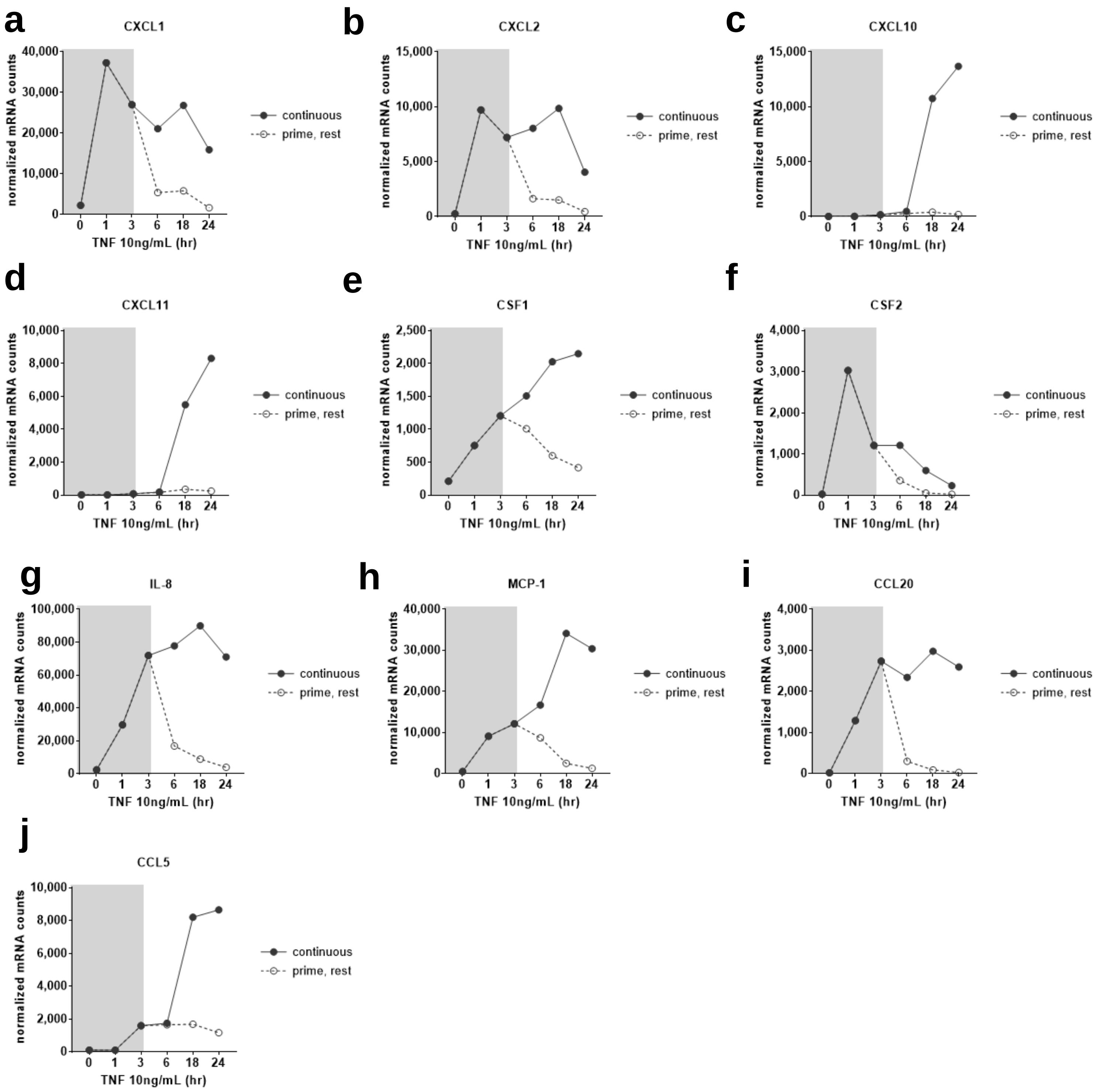


**Supplemental Figure 4.** Persistence of chemokine mRNA in HMEC-1 after 3hr priming with TNFα, followed by cytokine withdrawal for 3hr, 15hr or 21hr.

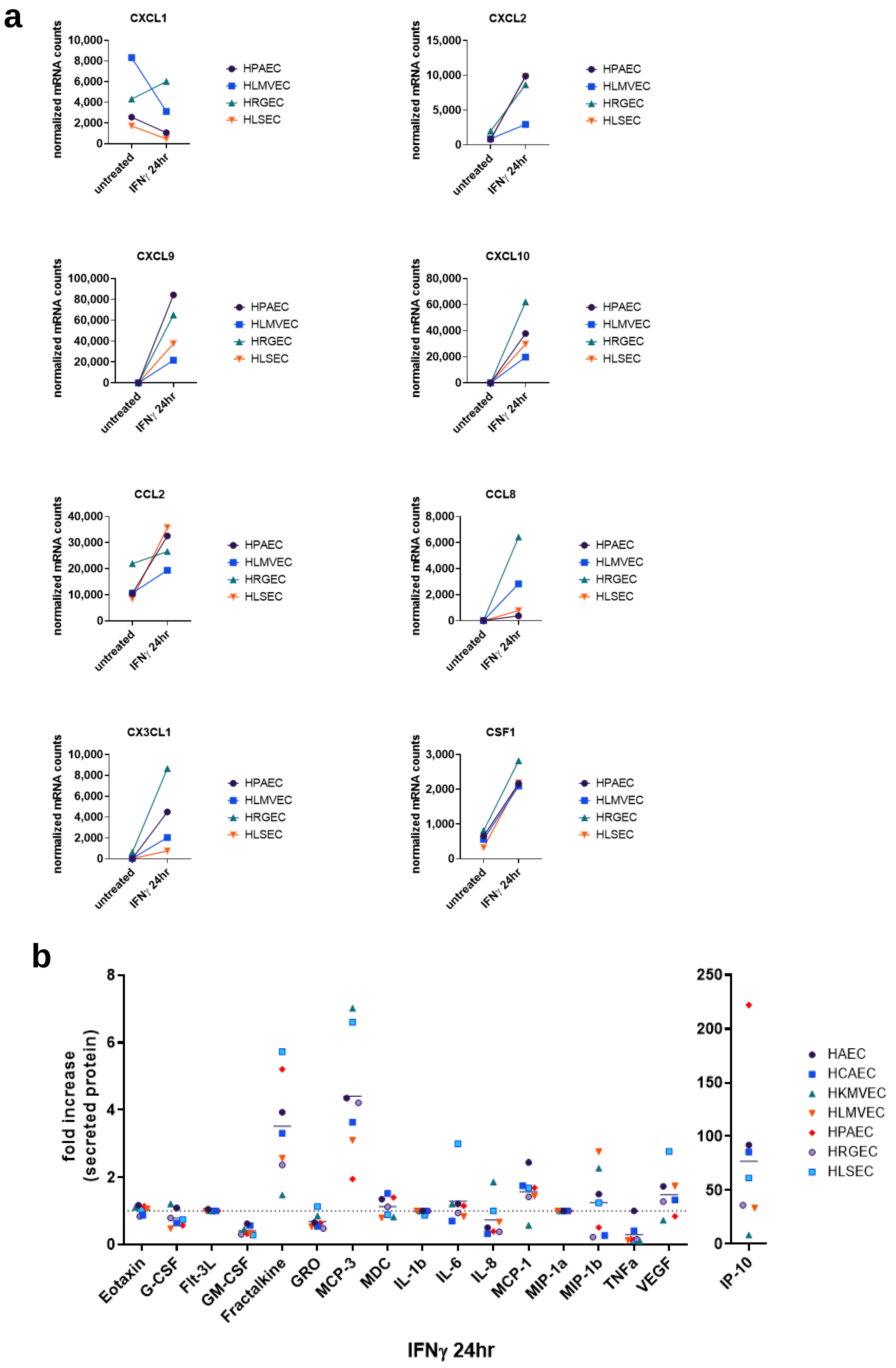
a
CXCL1
normalized mRNA counts
10,000
8,000
6,000
4,000
2,000
0
untreated
IFNγ 24hr
HPAEC
HLMVEC
HRGEC
HLSEC
CXCL2
15,000
10,000
5,000
CXCL9
100,000
80,000
60,000
40,000
20,000
CXCL10
CCL2
40,000
30,000
20,000
CCL8
CX3CL1
CSF1
3,000
2,000
1,000
b
fold increase (secreted protein)
Eotaxin
G-CSF
Flt-3L
GM-CSF
Fractalkine
GRO
MCP-3
MDC
IL-1b
IL-6
IL-8
MCP-1
MIP-1a
MIP-1b
TNFa
VEGF
IP-10
250
200
150
100
50
HAEC
HCAEC
HKMVEC
HLMVEC
HPAEC
HRGEC
HLSEC
IFNγ 24hr

**Supplemental Figure 5. IFNγ induced chemokine expression across primary endothelial vascular beds.**

Primary human pulmonary artery, lung microvascular, renal glomerular and liver microvascular endothelial cells were stimulated with IFNγ (200U/mL) for 24hr. a) mRNA counts of chemokine genes were measured by Nanostring. b) Secreted chemokine protein was measured by Luminex.

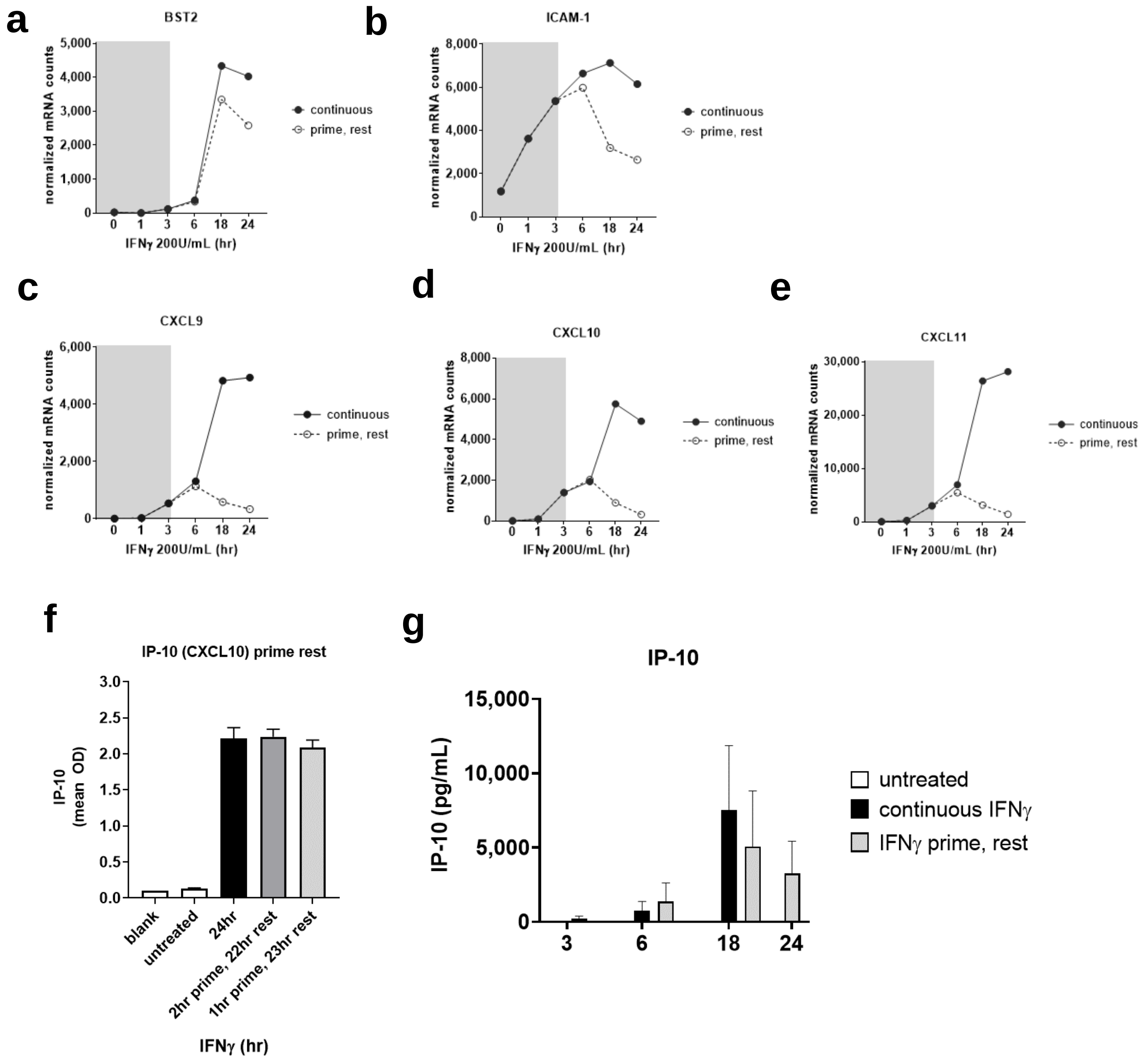


**Supplemental Figure 6.** Persistence of mRNA in HMEC-1 after 3hr IFNγ priming and withdrawal for an additional 3hr, 15hr or 21hr.
**a-e.** Normalized mRNA counts are graphed from one representative experiment with HMEC-1.
**f-g.** Secreted IP-10 was measured by ELISA, n=3-4 independent experiments.

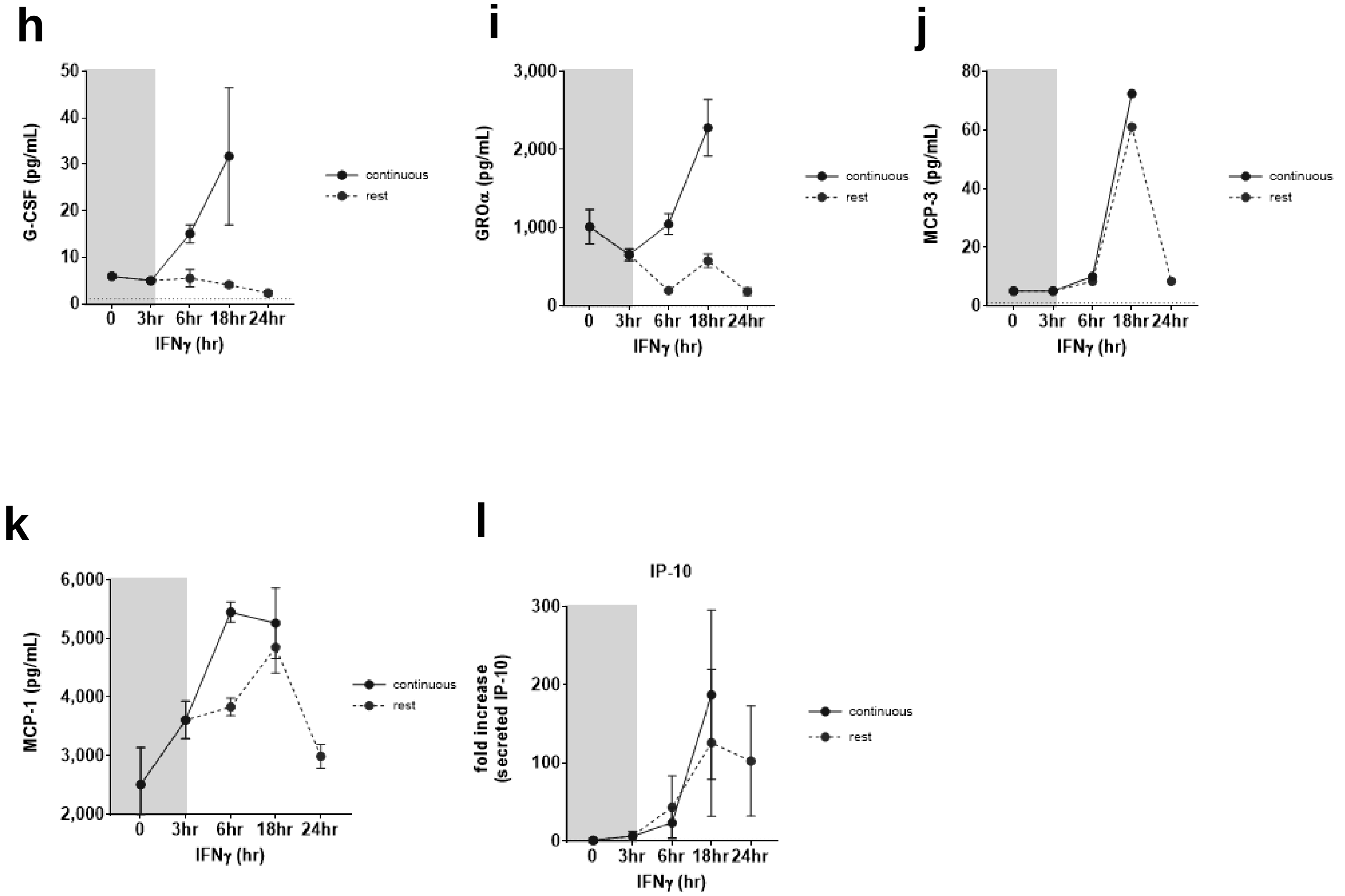


**Supplemental Figure 6.** Persistence of mRNA in HMEC-1 after 3hr IFNγ priming and withdrawal for an additional 3hr, 15hr or 21hr.

**h-l** Secreted chemokines were measured by Luminex (n=3).

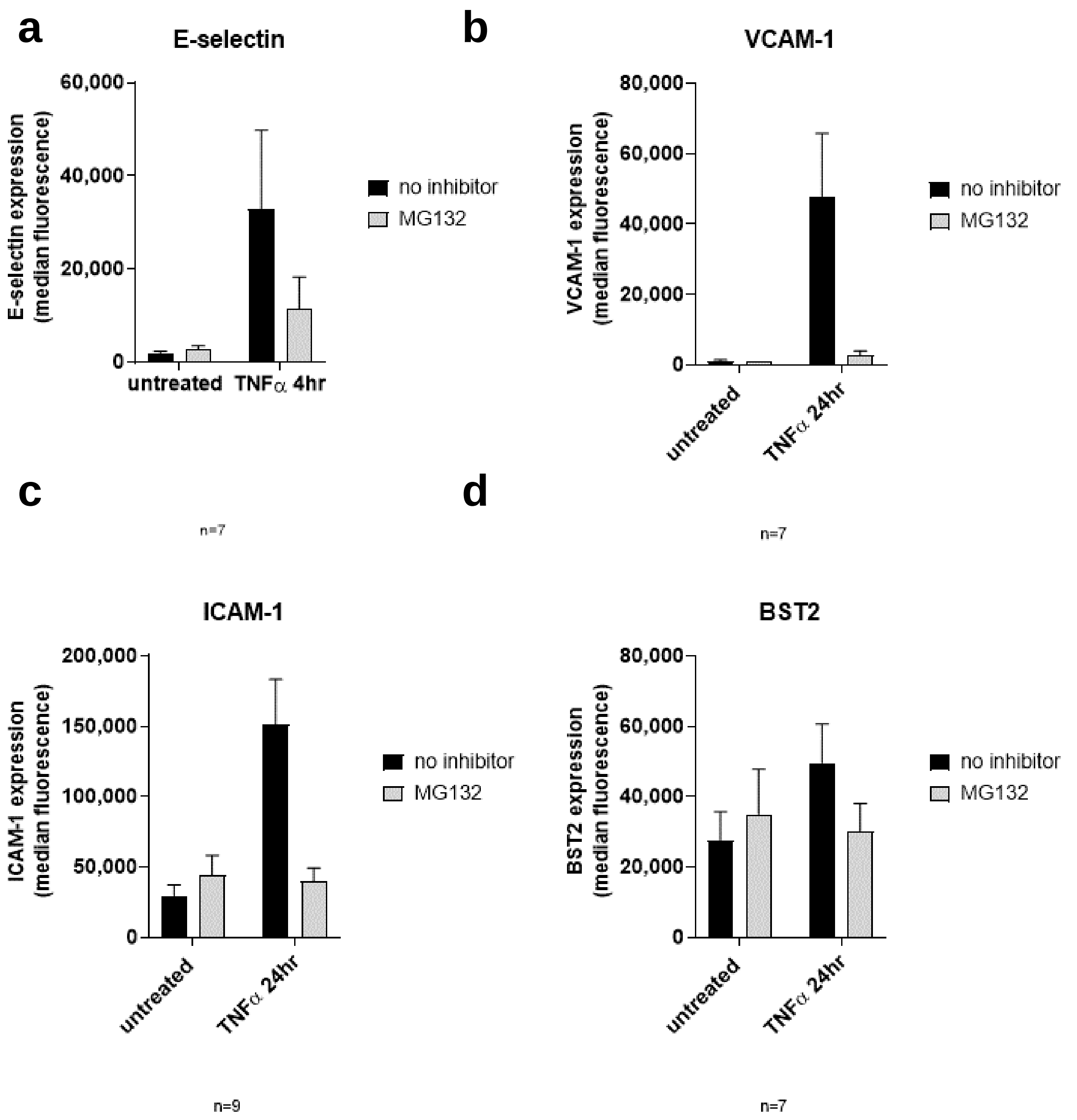


**Supplemental Figure 7.** NFκB dependence of TNFα induced adhesion molecules. HAEC were left uninhibited or pre-treated with the NFκB inhibitor MG132 (5μM) for 30min, prior to continuous stimulation with TNFα. Cell surface adhesion molecules were measured by flow cytometry. Raw MFI values for E-selectin (a), VCAM-1 (b), ICAM-1 (c), and BST2 (d) are shown, averaged over 7-9 independent experiments.

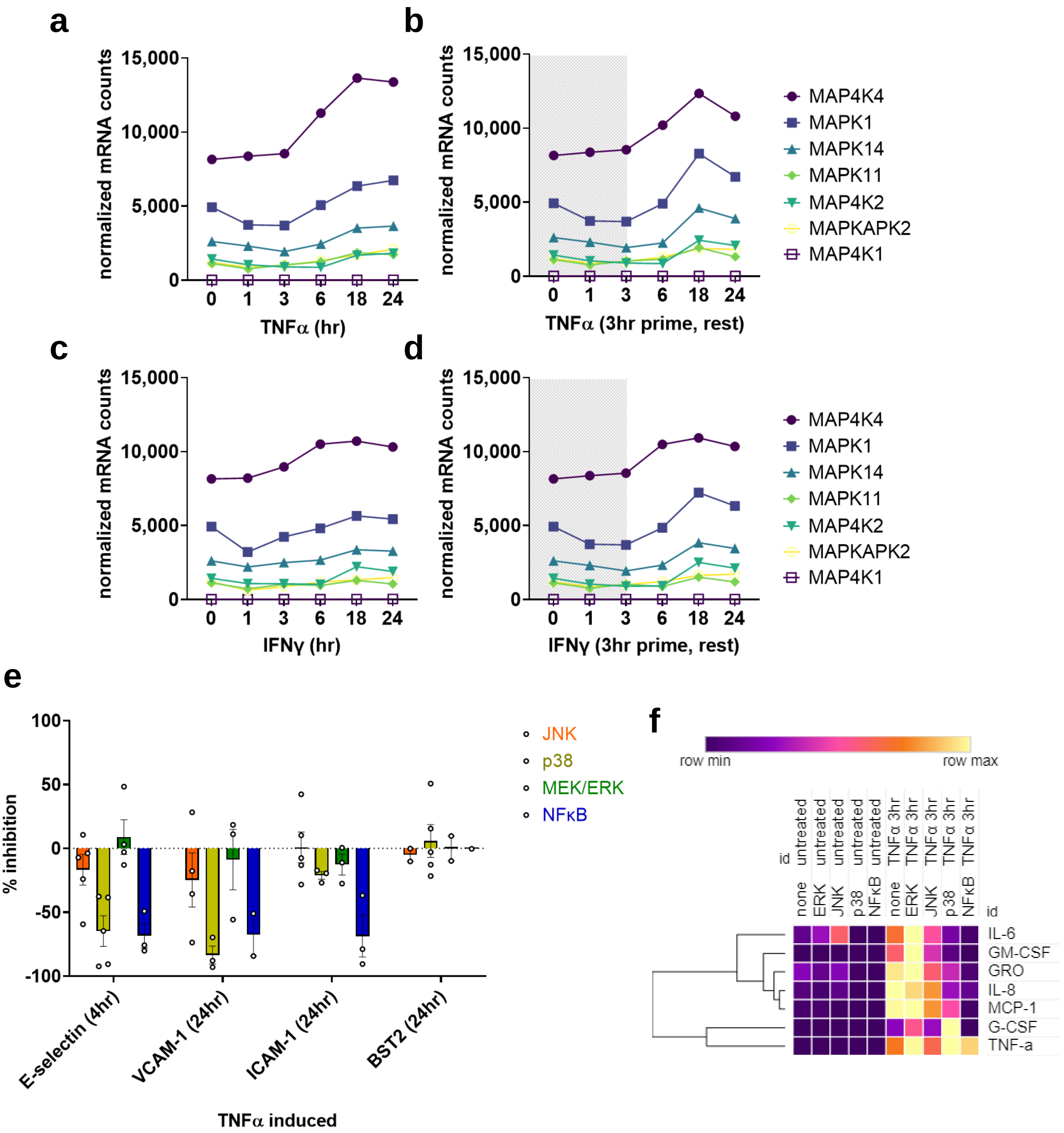


**Supplemental Figure 8.** Cooperative MAPK signaling in TNFα-mediated endothelial activation. a-d) HMEC-1 were stimulated with TNFα or IFNγ, and MAPK gene expression was measured by Nanostring. e, f) HAEC were pre-treated with the JNK inhibitor (SP600125, 20μM), p38 inhibitor (SB203580, 25μM) or MEK/ERK inhibitor (U0126, 10μM) prior to stimulation with TNFα. Cell surface adhesion molecules were measured by flow cytometry (e) and percent inhibition of expression is presented. Secreted chemokines were measured by Luminex (f) and relative protein concentrations are shown in the heat map.

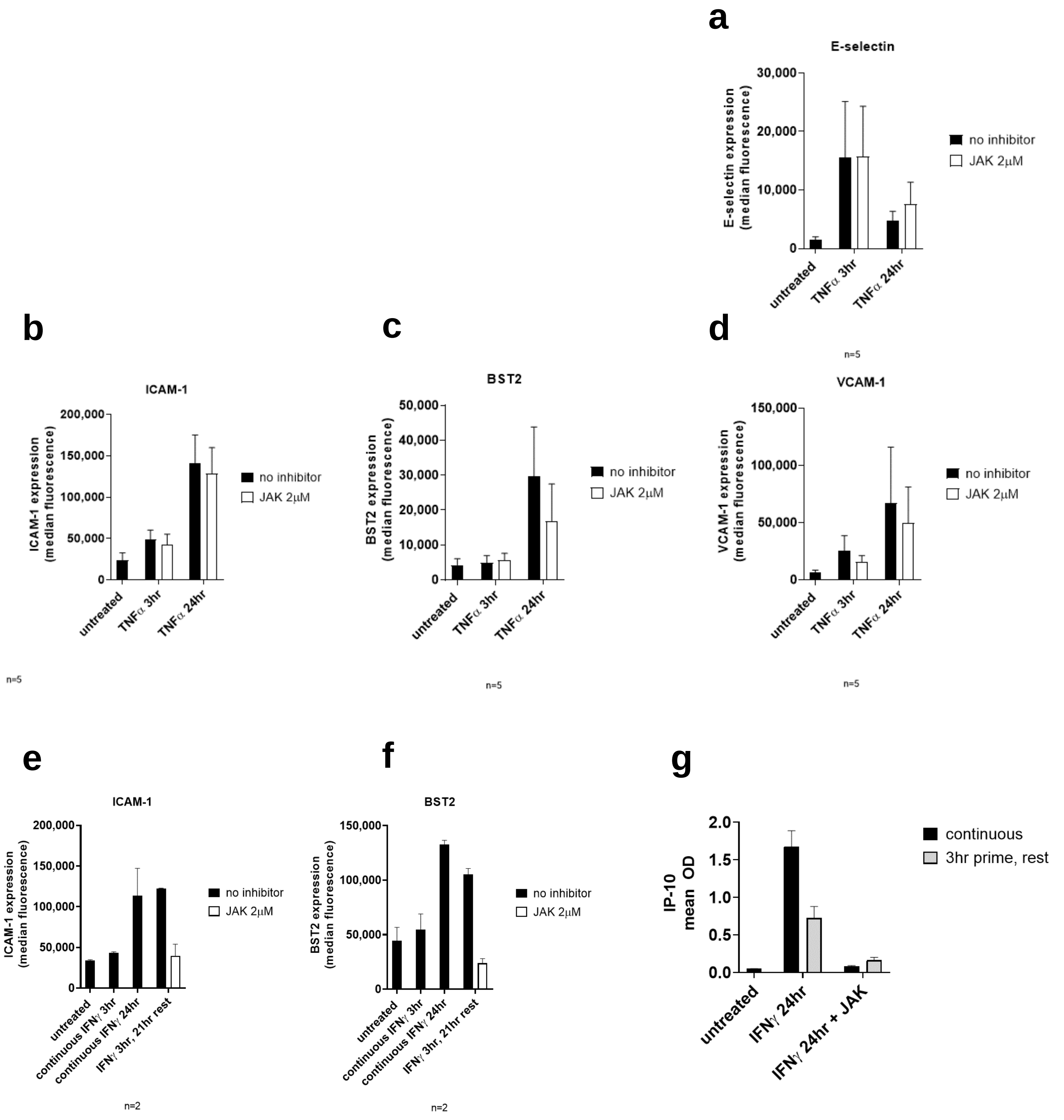


**Supplemental Figure 9.** JAK dependence of TNFα induced adhesion molecules. HAEC were left uninhibited or pre-treated with the JAK1/2 inhibitor ruxolitinib (2µM) for 30min, prior to continuous stimulation with TNFα or IFNγ. Cell surface adhesion molecules were measured by flow cytometry. Raw MFI values are shown, averaged over 2-5 independent experiments.

**h**

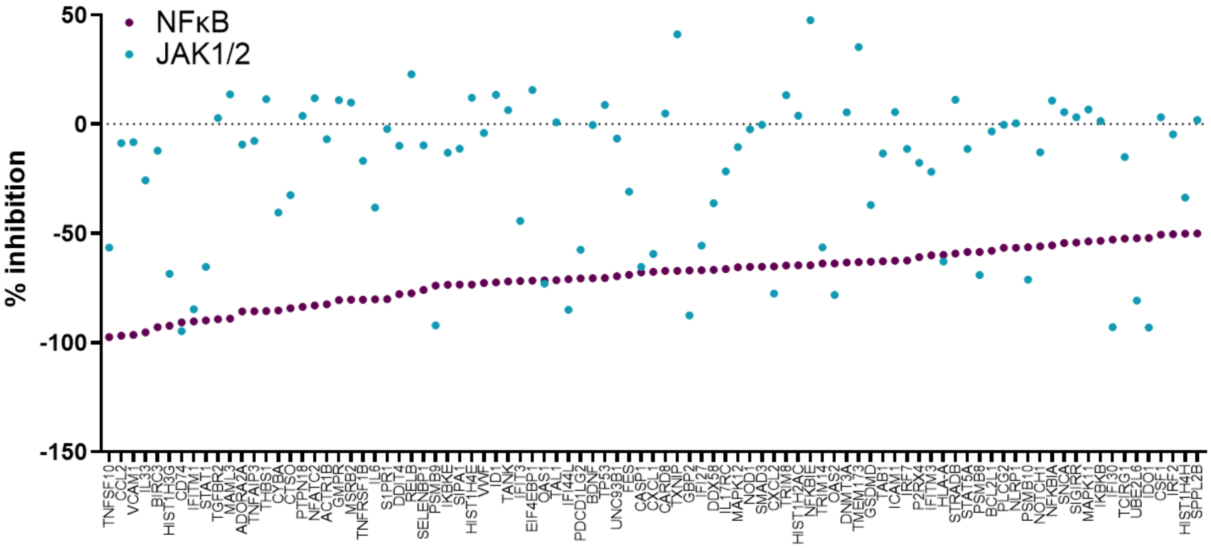


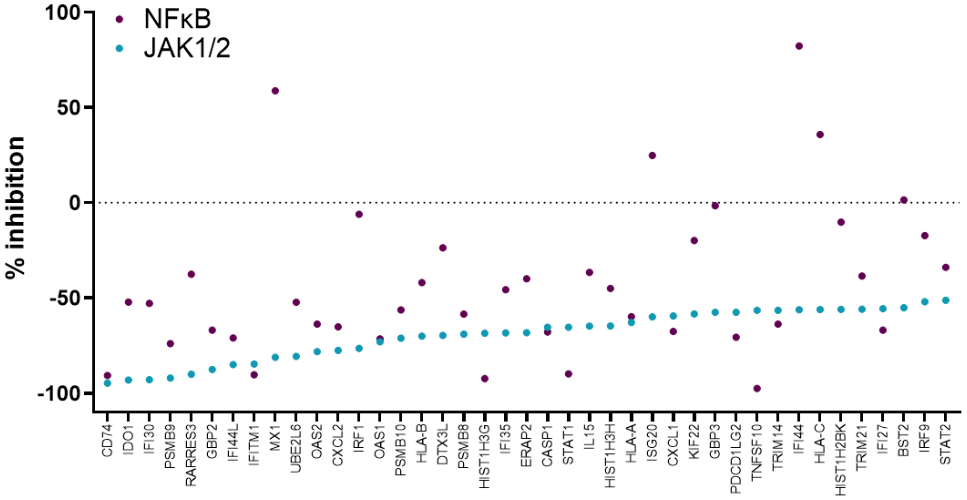


**i**

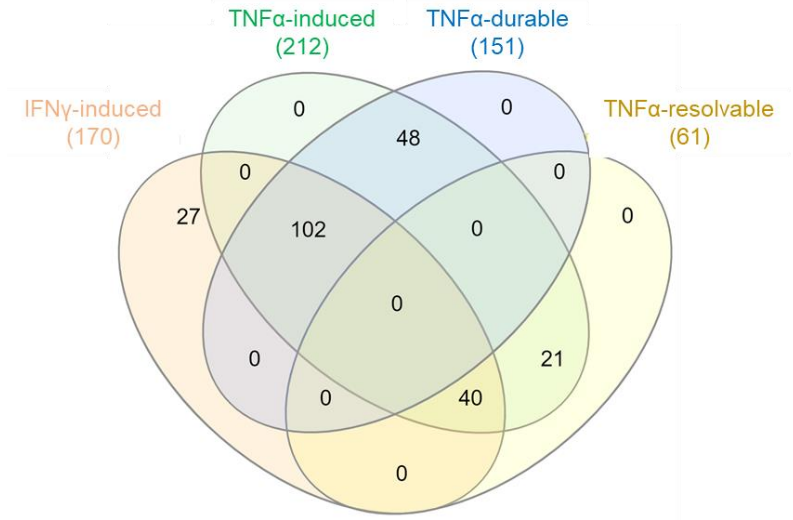


**Supplemental Figure 9 (continued).** JAK dependence of TNFα induced adhesion molecules. HAEC were left uninhibited or pre-treated with the JAK1/2 inhibitor ruxolitinib (2μM) or NFκB inhibitor MG132 (10μM) for 30min, prior to continuous stimulation with TNFα for 18hr. The percent change in TNFα-induced genes (≥1.5-fold) is shown.

j

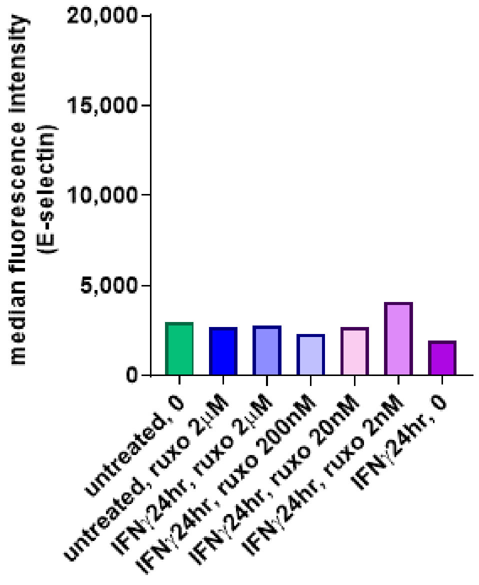


k

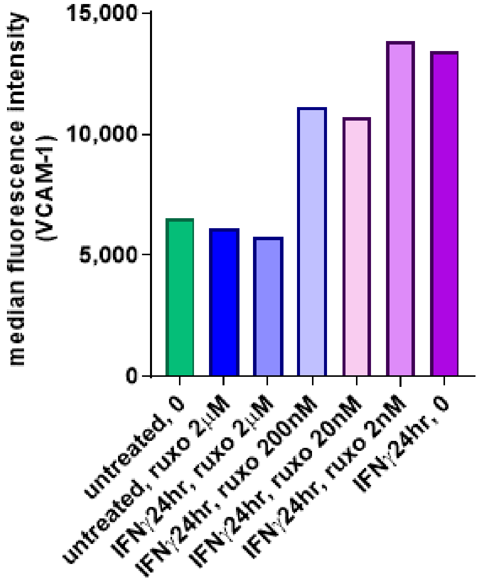


l

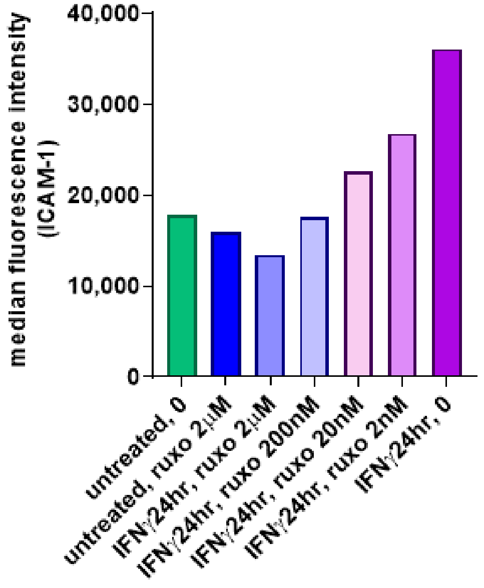


m

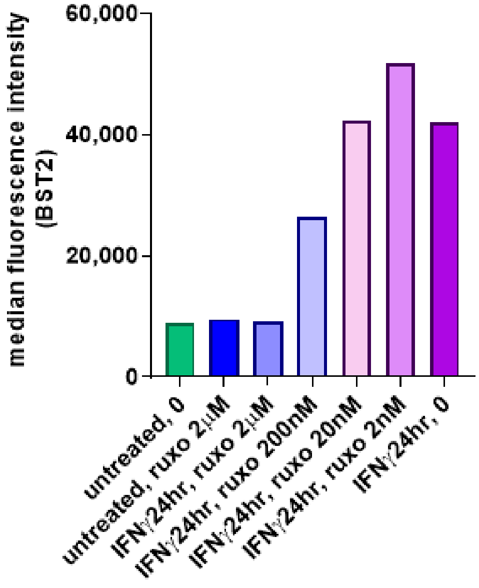


**Supplemental Figure 9 (continued).** HAEC were left uninhibited or pre-treated with 10-fold serial dilutions of the JAK1/2 inhibitor ruxolitinib (0.02-2μM) for 30min, prior to continuous stimulation with IFNγ. Cell surface adhesion molecules were measured by flow cytometry. Raw MFI values are shown, representative of 2 independent preliminary experiments.
